# Simultaneous profiling of native-state proteomes and transcriptomes of neural cell types using proximity labeling

**DOI:** 10.1101/2025.01.29.635500

**Authors:** Christina C. Ramelow, Eric B. Dammer, Hailian Xiao, Lihong Cheng, Prateek Kumar, Claudia Espinosa-Garcia, Maureen M. Sampson, Ruth S. Nelson, Sneha Malepati, Dilpreet Kour, Rashmi Kumari, Qi Guo, Pritha Bagchi, Duc M. Duong, Nicholas T. Seyfried, Steven A. Sloan, Srikant Rangaraju

## Abstract

Deep molecular phenotyping of cells at transcriptomic and proteomic levels is an essential first step to understanding cellular contributions to development, aging, injury, and disease. Since proteome and transcriptome level abundances only modestly correlate with each other, complementary profiling of both is needed. We report a novel method called simultaneous protein and RNA –omics (SPARO) to capture the cell type-specific transcriptome and proteome simultaneously from both in vitro and in vivo experimental model systems. This method leverages the ability of biotin ligase, TurboID, to biotinylate cytosolic proteins including ribosomal and RNA-binding proteins, which allows enrichment of biotinylated proteins for proteomics as well as protein-associated RNA for transcriptomics. We validated this approach first using well-controlled in vitro systems to verify that the proteomes and transcriptomes obtained reflect the ground truth, bulk proteomes and transcriptomes. We also show that the effect of a biological stimulus (e.g., neuroinflammatory activation by lipopolysaccharide) can be faithfully captured. We also applied this approach to obtain native-state proteomes and transcriptomes from two key neural cell types, astrocytes and neurons, thereby validating the in vivo application of SPARO. Next, we used these data to interrogate protein-mRNA concordance and discordance across these cell types, providing insights into groups of molecular processes that exhibit uniform or cell type-specific patterns of mRNA-protein discordance.

## Introduction

The transcriptome and proteome of a cell provide complementary molecular information about its phenotype, state, and cellular functions. Quantifying RNA and protein levels in cell types using high-throughput –omics approaches uncover mechanisms and molecular pathways that contribute to development, aging, injury, and disease. RNA-sequencing can be leveraged to record different transcript species, expression patterns, gene structure, splicing patterns, and post-transcriptional modifications. On the other hand, mass spectrometry (MS)-based quantitative proteomics provides information on protein abundances, localization, post-translational modifications, structure, cell-to-cell interactions, and effector functions. Importantly, the number of RNA transcripts for a given gene does not always correlate with its protein abundance^1,2^. This is straightforward to explain from a biological standpoint by considering dynamics of RNA stability, translational regulation, protein half-life, chaperone proteins, cellular sequestration, as well as other post-transcriptional and post-translational events^3–7^. This contributes to observed discordances between mRNA and protein abundances and molecular signatures in cells and tissues.

Measuring RNA and protein levels simultaneously can reveal changes at the RNA-level that are not reflected at the protein-level, and vice-versa, and enables exploration of mechanisms regulating mRNA-protein correlation. This can have biological implications as well as highly practical consequences that impact choice of RNA or protein markers in experiments (e.g., choice of RNA probe for in situ hybridization or choice of protein marker for spatial studies). Also, the level of discordance between mRNA and protein, and the mechanisms that control mRNA and protein levels may vary across different cell types, tissues, and biological contexts, and cannot be ascertained from bulk tissue – omics approaches. Therefore, we need new approaches to jointly measure mRNA and protein levels in cell type-specific contexts.

Several attempts have been made to quantify this general concordance (or discordance) between transcriptomic and proteomic levels^2,8–16^. Simultaneous sampling of the transcriptome and proteome can be readily accomplished under in vitro monoculture conditions. However, under in vivo settings, these methods involve collecting RNA and protein from separate physical samples and rely on cell type purification using mechanical approaches, which, by themselves, impact transcriptomic and molecular characteristics of the cells^17^. The ability to purify intact cells from complex tissues like the brain also varies across cell type. For example, adult neurons are difficult to isolate without loss of their cellular and synaptic architecture^18,19^.

A general approach to obtain cell type-specific transcriptomes or proteomes from tissues that are independent of cell type isolation involves protein tagging. For cell type-specific in vivo transcriptomics, ribosomal subunits (e.g., Rpl22) can be tagged (e.g., HA) in a cell type-specific manner using Cre/lox genetics, allowing capture of ribosome-associated mRNA species (translatome). For cell type-specific proteomics, metabolic labeling with non-canonical amino acids^20–25^ and proximity-labeling by biotin ligases^26–37^ have been employed. There have also been several efforts to achieve joint mRNA and protein measurements, using RiboTag^38–41^ and Translating Ribosome Affinity Purification (TRAP)^42,43^ to capture mRNAs and nascent peptides from a single tagged ribosomal protein under a cell type-specific promoter. However, these approaches limit profiling to only ribosome-bound transcripts and nascent peptides rather than the broader proteome and transcriptome of the cell. Further, mammalian cells contain at least 79 different ribosomal proteins for cytosolic translation^44^, and ribosomes with different protein compositions translate different groups of mRNAs^45^. Therefore, ribosome affinity purification-based methods that target a single ribosomal protein may not capture the full complexity of the RNA species within a cell.

Another method to achieve joint mRNA and protein measurements is Single-Cell Protein And RNA Co-profiling (SPARC)^46^, which uses poly(A) oligo-dTs conjugated to magnetic beads that hybridize to mRNAs in conjunction with a homogeneous protein extension assay (PEA)^47,48^. SPARC, however, is limited by the 96-plex capability of the targeted proteins of the PEA assay. Due to the limitations of current methods, new tools for complementary molecular profiling of cell type-specific transcriptomes and proteomes are needed to gain a more global characterization of cell types in their native-states in a complex tissue environment.

To develop an approach for **s**imultaneous **p**rotein and **R**NA **o**mics (SPARO), we leveraged the biotin ligase, TurboID. Using MS-based proteomics, we previously found that cytosolic TurboID biotinylates many RNA-binding proteins (RBPs) and ribosomal proteins both in vitro and in vivo, including Rpl22 that is used in the RiboTag approach^34,35^. Therefore, we hypothesized that transcripts associated with these proteins could also be purified simultaneously with biotinylated proteins. We developed and validated the SPARO approach to capture cell type-specific transcriptomes and proteomes using a microglial cell line in vitro, and neurons and astrocytes in vivo. This allowed us to investigate the concordance and discordance between mRNA and protein levels, and to examine how patterns of mRNA-protein discordance are conserved or vary across cell types. SPARO represents a novel approach to simultaneously quantify RNA and protein profiles with potentials for broad applications in vitro and in vivo.

## Results

### Validation of SPARO in BV2-TurboID cells in vitro

To test whether we could leverage TurboID-based cytosolic proteomic labeling to simultaneously capture cellular proteomes and transcriptomes, we first used our previously validated, stably transduced mouse microglia BV2 cell line that expresses V5-TurboID-NES (BV2-TurboID)^35,49^. In this construct, the TurboID biotin ligase is restricted to the extranuclear compartment via a nuclear export sequence (NES) to bias towards biotinylation of the cytosolic proteome (**Fig. 1a**). By starting with a homogeneous cell line, we could quantitatively assess how the transcriptome and proteome of BV2 cells using the SPARO approach compare with wholecell transcriptomes and proteomes from bulk cell lysates. Furthermore, to test whether SPARO could also capture cellular changes in microglia induced by an inflammatory stimulus, we treated BV2 control and BV2-TurboID cells with lipopolysaccharide (LPS) (**Fig. 1b**). We lysed cells in a buffer that maintains RNA-protein interactions and then processed for simultaneous transcriptomics (mRNA-seq) and proteomics (label-free quantitative mass spectrometry or LFQ-MS). Immunoblot analyses confirmed the presence of biotinylated proteins (via streptavidin) and the TurboID protein (via V5) in BV2-TurboID cells, which were absent in non-TurboID control cells both before (**Supplementary Fig. 1a**) and after streptavidin bead affinity purification (henceforth “pulldown”) (**Fig. 1c**). We detected only endogenously biotinylated proteins in the BV2 control cells (**Fig. 1c**). After streptavidin pulldown, we confirmed protein enrichment using a silver stain (**Supplementary Fig. 1b**). RNA gel electrophoresis showed comparably high levels of RNA in bulk BV2 lysates (**Supplementary Fig. 1c**). Importantly, we found high levels of RNA in BV2-TurboID pulldowns but negligible RNA in BV2 control cell pulldowns without TurboID (**Fig. 1d**), confirming that RNA is enriched only when proteomic biotinylation occurs in BV2 microglia.

**Figure 1:**
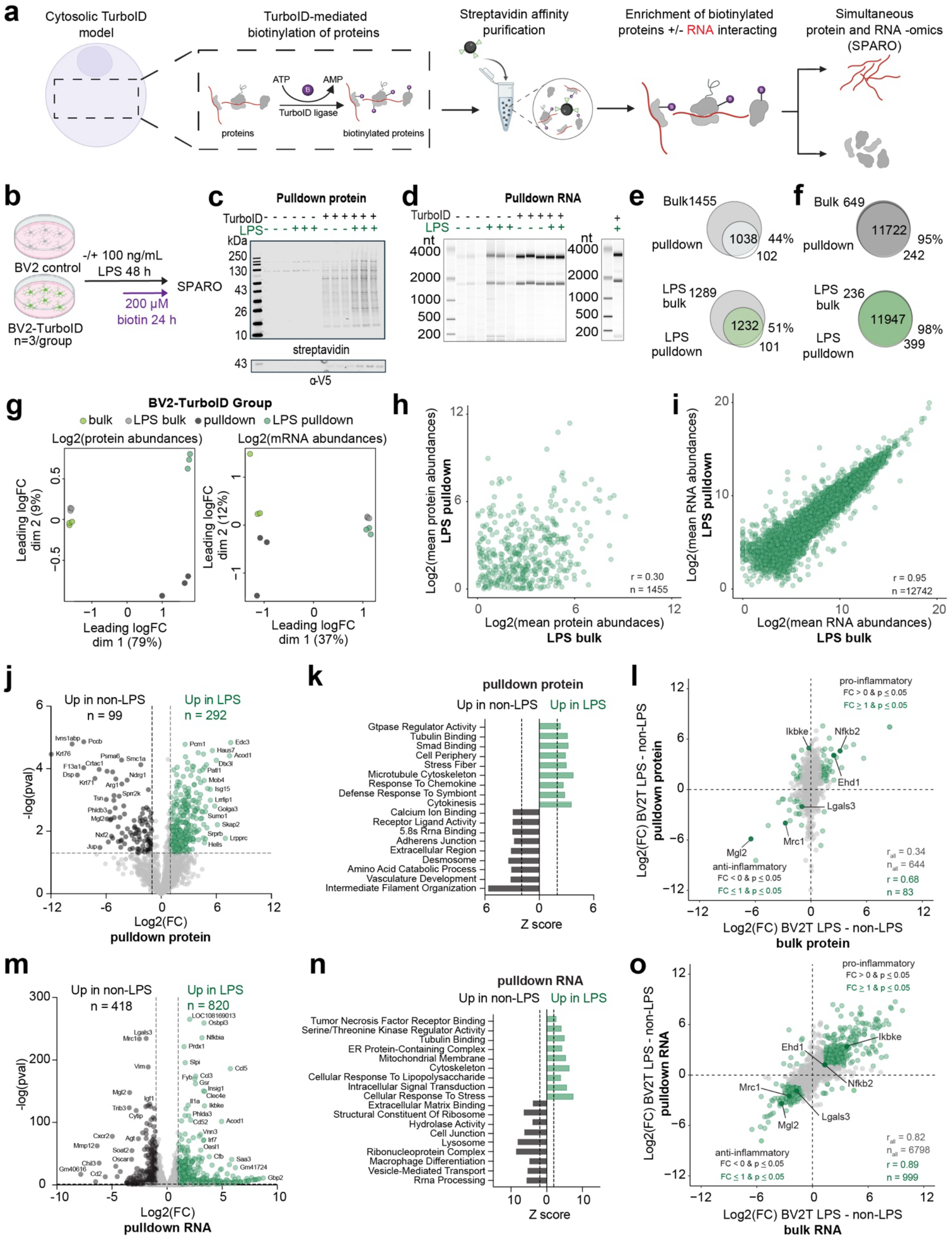
Simultaneous transcriptomics and proteomics profiling of BV2 microglia under homeostatic and LPS-stimulated conditions. **a**. Schematic for leveraging cytosolic TurboID to create a simultaneous protein and RNA –omics (SPARO) approach. **b**. Experimental design: BV2 control and BV2-TurboID cells (n = 3 biological replicates/genotype/condition) were treated with 100 ng/mL of LPS for 48 hours and 200 μM of biotin for 24 hours during the second day of LPS treatment. **c**. Immunoblot visualization of biotinylated proteins from BV2-TurboID cells at the pulldown level not seen in non-TurboID control cells (probed with streptavidin-680). Presence of TurboID recombinant protein (via V5 tag) was also detected at the pulldown level. **d**. RNA gel electrophoresis of RNA eluted off biotinylated proteins after streptavidin pulldown. High levels of rRNA and mRNA were detected in BV2-TurboID cells, but not seen in non-TurboID control cells. **e**. Venn diagram representation of protein numbers identified by LFQ-MS present in the BV2-TurboID bulk and pulldown proteomes without LPS treatment (top) and with LPS treatment (bottom). **f**. Venn diagram representation of RNA quantities identified by RNA-seq present in the BV2-TurboID bulk and pulldown transcriptomes without LPS treatment (top) and with LPS treatment (bottom). **g**. MDS plot of normalized LFQ-MS intensity data (left) in each BV2-TurboID proteome group. **h**. Correlation analysis of protein abundances from normalized LFQ-MS mean intensity values between the BV2-TurboID bulk and pulldown proteomes (n = 1455 proteins, Pearson correlation coefficient r = 0.30). **i**. Correlation analysis of RNA abundances from normalized RNA-seq mean count values between the BV2-TurboID bulk and pulldown transcriptomes (n = 12742 genes, Pearson correlation coefficient r = 0.95). **j**. Volcano plot representation of DEPs between non-LPS and LPS-treated BV2-TurboID cells at the pulldown level (n=3/group). Black symbols (two-sided t-test, p ≤ 0.05 and ≥ 1-fold change, n = 99) represent DEPs in non-LPS-treated BV2-TurboID cells. Green symbols (two-sided t-test, p ≤ 0.05 and ≥ 1-fold change, n = 292) represent DEPs in LPS-treated BV2-TurboID cells. **k**. Proteome GO analysis of DEPs visualized in panel j, showing overrepresented ontologies in non-LPS treated (black) and LPS-treated (green) BV2-TurboID cells at the pulldown level. **l**. Correlation analysis of all shared DEPs (two-sided t-test, p ≤ 0.05) between the LPS-treated BV2-TurboID cells at the bulk and pulldown level (n = 644 proteins, Pearson correlation coefficient r = 0.34). Green symbols represent highly enriched shared DEPs (two-sided t-test, p ≤ 0.05 and ≥ 1-fold change, n = 83, Pearson correlation coefficient r = 0.68). Examples of pro-inflammatory and anti-inflammatory canonical microglia markers labeled and in bold. **m**. Volcano plot representation of differentially expressed genes (DEGs) between non-LPS and LPS-treated BV2-TurboID cells at the pulldown level (n=3/group). Black symbols (two-sided t-test, p ≤ 0.05 and ≥ 1-fold change, n = 418) represent DEGs enriched in non-LPS-treated BV2-TurboID cells. Green symbols (two-sided t-test, p ≤ 0.05 and ≥ 1-fold change, n = 820) represent DEGs enriched in LPS-treated BV2-TurboID cells. **n**. Transcriptome GO analysis of DEGs visualized in panel m. of overrepresented ontologies of non-LPS treated black) and LPS-treated BV2-TurboID cells (green scale) at the pulldown level. **o**. Correlation analysis of all shared DEGs (two-sided t-test, p ≤ 0.05) between the LPS-treated BV2-TurboID cells at the bulk and pulldown level (n = 6798 genes, Pearson correlation coefficient r = 0.82). Green symbols represent highly enriched shared DEGs (two-sided t-test, p ≤ 0.05 and ≥ 1-fold change, n = 999, Pearson correlation coefficient r = 0.89). Examples of pro-inflammatory and anti-inflammatory canonical microglia markers are labeled and in bold.

We next performed LFQ-MS and RNA-seq on bulk BV2-TurboID samples and pulldowns. Raw intensity and processed intensity values are found in Supplementary Data 2 and 3, respectively. Raw RNA-seq and processed counts values are found in Supplementary Data 4 and 5, respectively. We first assessed the overlap of the number of proteins between the TurboID pulldowns and the bulk proteome. We observed 44% (untreated) and 51% (LPS-treated) overlap of TurboID pulldown proteomes with the ground truth bulk BV2 proteomes, fairly consistent with prior observations in BV2-TurboID cells^35^. This is likely a result of the NES tag on TurboID that biases the proteome towards the cytosolic compartment and against the nuclear and mitochondrial compartments (**Fig. 1e**). Next, to benchmark the fidelity of our TurboID pulldown transcriptome, we compared our RNA-seq datasets between the bulk BV2 samples and our pulldown populations. We observed between 95% (untreated) and 98% (LPS-treated) overlap of TurboID pulldown transcriptomes with the ground truth bulk BV2 transcriptome (**Fig. 1f**). We next asked whether the pulldown proteomes and transcriptomes could adequately capture changes in BV2 cells upon exposure to LPS (**Fig. 1g, left**). Multidimensional scaling (MDS) analysis of the transcriptomes identified distinct untreated and LPS-treated transcriptomes regardless of sample type (bulk vs pulldown) (**Fig. 1g, right**) suggesting that LPS-effect was equally captured by the TurboID pulldown and bulk transcriptomes.

To test the fidelity of the pulldowns to their respective bulk samples, we next performed correlation analysis of our LFQ-MS and RNA-seq data, respectively. At the proteomic level, we observed modest correlation between LPS-treated pulldown and the bulk LPS-treated BV2-TurboID proteomes (Pearson correlation coefficient r = 0.30, n = 1455 shared proteins, **Fig. 1h**). Notably, we observed a much higher correlation between the LPS-treated pulldown transcriptomes and bulk BV2-TurboID transcriptomes (Pearson correlation coefficient r = 0.95, n = 12742 shared transcripts, **Fig. 1i**). We also compared the correlation between the untreated bulk and pulldown proteomes and transcriptomes, respectively (**Supplementary Fig. 1e and h**). To confirm the cytosolic bias of the V5-TurboID-NES recombinant protein, we differentially compared the pulldown and bulk BV2-TurboID transcriptomes and proteomes. Differential enrichment analysis of BV2 transcriptomic and proteomic data are found in Supplementary Data 6 and 7, respectively. At the transcriptomic level, we found that mitochondrial-encoded genes (e.g., *COX1, CYTB* and *ND2*) were highly enriched in the bulk samples (p ≤ 0.05, ≥ 1-fold change, **Supplementary Fig. 2a**) as compared to the pulldowns, consistent with inability of cytosolic TurboID to label intra-mitochondrial proteins and their associated transcripts. At the proteomic level, the bulk proteome was enriched in nuclear and mRNA-splicing related terms, consistent with under-sampling of the nuclear proteome by TurboID labeling (p ≤ 0.05, ≥ 1-fold change, **Supplementary Fig. 3a**). SPARO also detected proteomic and transcriptomic changes driven by LPS treatment (**Fig. 1j and m, Supplementary Fig. 2 and 3**). As expected, GO analysis of differentially enriched proteins (DEPs) between TurboID pulldowns revealed overrepresented ontologies of LPS-treated cells including response to chemokine, defense response to symbiont and cytokinesis (**Fig. 1k, Supplementary Data 8**). The changes that we observed upon LPS treatment measured by TurboID pulldown were still modestly correlated with LPS changes in the bulk proteome (Pearson correlation coefficient r = 0.34, n = 644 shared proteins, p ≤ 0.05, **Fig. 1l**). However, this correlation was substantially improved when specifically considering proteins that exhibit high differential abundance between LPS-treated and control BV2-TurboID cells (Pearson correlation coefficient r = 0.83, n = 83 shared proteins with p ≤ 0.05 and ≤ –1 or ≥ 1-fold change, **Fig. 1l**). We also found canonical pro-inflammatory microglial activation markers (e.g., Ikbke, Nfkb2, and Ehd1) enriched in the LPS-treated BV2-TurboID pulldowns and anti-inflammatory microglial activation markers (e.g., Mrc1/Cd206, Mgl2 and Lgals3) enriched in the control pulldowns (p ≤ 0.05, ≥ 1-fold change, **Fig.1l**).

Similarly to the proteome, we performed differential enrichment and GO analyses of differentially expressed genes (DEGs) in the LPS-treated and non-LPS treated pulldown transcriptomes (**Supplementary Data 9**). We observed the same canonical reactive microglial response enriched in LPS treated and untreated samples (**Fig. 1m-o**). Unlike at the protein level, the LPS-driven transcriptomic changes correlated highly between the TurboID RNA pulldown and the LPS-enriched bulk transcriptome (Pearson correlation coefficient r = 0.82, n = 6798 shared transcripts with p ≤ 0.05, **Fig. 1o**). This correlation was even more improved when specifically considering genes that exhibit high differential enrichment between LPS-treated and control BV2-TurboID cells (Pearson correlation coefficient r = 0.89, n = 999 shared genes, p ≤ 0.05 and ≤ –1 or ≥ 1-fold change, **Fig. 1o**). Additional transcriptomic and proteomic GO analysis data comparing either LPS-treated versus untreated bulk or pulldown samples and pulldown versus bulk samples can be found in Supplemental Data 10-15.

Overall, these results suggest that SPARO can simultaneously capture the cellular transcriptome and proteome and effectively captures inflammatory effects of LPS in an in vitro mammalian system.

### Validation of SPARO in vivo using cortical astrocytes and neurons

We next wanted to test whether we could capture cell type-specific transcriptomes and proteomes simultaneously in vivo while retaining the native state of these cells. We crossed Rosa26^TurboID/wt^ (TurboID) mice with appropriate inducible Cre-ERT2 mice, to drive Cre expression specifically in Camk2a-expressing excitatory neurons or in Aldh1l1-expressing astrocytes. We chose the Aldh1l1-Cre/ERT2 mice and Camk2a-Cre/ERT2 mouse lines for their extensive validation and non-leakiness of Cre^40,50^. Our breeding scheme resulted in the generation of Aldh1l1^CreERT2/wt^/Rosa26^TurboID/wt^ (astrocyte-TurboID) and Camk2a^CreERT2/wt^/Rosa26^TurboID/wt^ (neuron-TurboID) mice. We used Aldh1l1^CreERT2/wt^ or Camk2a^CreERT2/wt^ as Cre-only controls. We induced Cre-mediated recombination by intraperitoneal tamoxifen at 7 weeks of age, followed by a 3-week gap, and then biotin water supplementation for 2 weeks (**Fig. 2a**). Based on our prior study, V5-TurboID-NES presence and biotin treatment does not have adverse effects on the mice and does not impact homeostatic cellular functions^29,31,34^. Following biotin treatment (age 2.5-3 months), we harvested and lysed cortical tissue in the same buffer as the in vitro studies to maintain RNA-protein interactions and then processed for simultaneous transcriptomics and proteomics using SPARO (**Fig. 2a**). Immunofluorescent imaging of the cortex showed robust biotinylation in TurboID brains compared to controls (**Fig. 2b and Supplementary Fig. 4f**). Moreover, the biotinylation signal co-localized with the astrocytic marker, S100β (in astrocyte-TurboID mice) and neuronal marker, β-III-tubulin (in neuron-TurboID mice) confirming cell targeting (**Fig. 2b**). Importantly, our prior work validated that the biotinylation signal present in astrocyte– and neuron-TurboID animals is cell type-specific via the absence of signal in other brain cell types, and without reactive gliosis^34^. Immunoblot analysis of cortical lysates at the bulk level (**Supplementary Fig. 4a and b**) and after streptavidin affinity purification displayed biotinylated proteins from astrocyte-TurboID and neuron-TurboID pulldowns not seen in non-TurboID controls (**Fig. 2c**). The presence of the TurboID recombinant protein via V5 presence was also only present in astrocyte-TurboID and neuron-TurboID pulldowns and not non-TurboID controls (**Fig. 2c**) providing biochemical verification of cell type-specificity.

**Figure 2:**
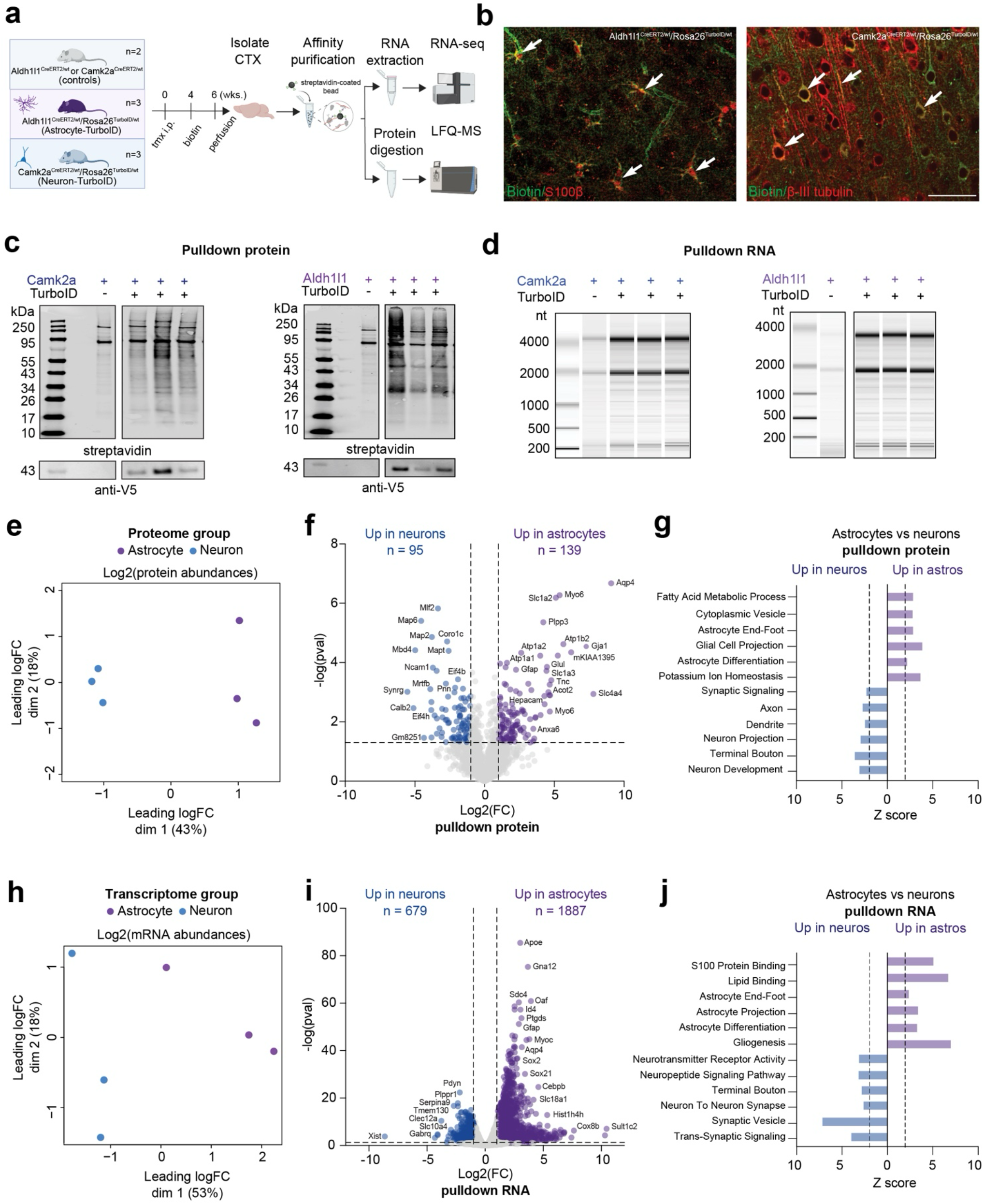
TurboID-based dual transcriptomic and proteomic analysis of cortical Aldh1l1-expressing astrocytes and Camk2a-expressing neurons in vivo. **a**. Experimental overview of dual affinity purification of RNA and protein using TurboID proximity labeling in Aldh1l1^CreERT2/wt^ or Camk2a^CreERT2/wt^ (control, n = 2 mice), Aldh1l1^CreERT2/wt^/Rosa26^TurboID/wt^ (astrocyte-TurboID, n = 3 mice) and Camk2a^CreERT2/wt^/Rosa26^TurboID/wt^ (neuron-TurboID, n = 3 mice) mouse models. Astrocyte-TurboID, neuron-TurboID and control mice received tamoxifen (75 mg/kg/day via intraperitoneal (i.p.) injections x 5 days). After 3 weeks of Cre recombination, mice received biotin-containing water (37.5 mg/L) for two weeks. Cortical tissue was isolated for streptavidin affinity purification, followed by LFQ-MS and RNA-sequencing. **b**. Representative immunofluorescence images from sagittal brain sections of control, astrocyte-TurboID and neuron-TurboID mice (n = 3 mice/experimental group) confirms astrocyte and neuron specific biotinylation via streptavidin-488 (green) overlapping with astrocytic marker S100β (red) and neuronal marker β-III-tubulin (red), respectively. Scale bar = 50 μm. **c**. Immunoblot visualization of cortical lysates after streptavidin affinity purification displaying biotinylated proteins from astrocyte-TurboID and neuronal-TurboID pulldowns not seen in non-TurboID controls (probed with streptavidin-680). The presence of the TurboID (via V5) was only present in astrocyte-TurboID and neuron-TurboID cortical pulldown tissue but not non-TurboID controls. **d**. RNA gel electrophoresis of RNA eluted off biotinylated proteins after streptavidin pulldown. High levels of rRNA and mRNA were detected in astrocyte-TurboID and neuronal-TurboID cortical tissue, not seen in non-TurboID controls. e. MDS plot of normalized LFQ-MS intensity data in astrocyte-TurboID and neuronal-TurboID pulldowns. **f**. Volcano plot representation of DEPs between astrocyte-TurboID and neuron-TurboID pulldowns (n=3/group). Blue symbols (two-sided t-test, p ≤ 0.05 and ≥ 1-fold change, n = 95) represent pulldown proteins enriched in neuron-TurboID cortical samples. Purple symbols (two-sided t-test, p ≤ 0.05 and ≥ 1-fold change, n = 139) represent pulldown proteins enriched in astrocyte-TurboID cortical samples. **g**. Proteome GO analysis of DEPs visualized in panel f. of overrepresented ontologies of neuron-TurboID (blue) and astrocyte-TurboID (purple) pulldowns, highlighting ontologies related to neuronal (e.g., synaptic signaling and neuron projection) and astrocyte function (fatty acid synthesis and glial cell projection), respectively. **h**. MDS analysis of normalized RNA-seq count data in astrocyte-TurboID and neuron-TurboID pulldowns. **i**. Volcano plot representation of DEGs between astrocyte-TurboID and neuron-TurboID pulldowns (n=3/group). Blue symbols (two-sided t-test, p ≤ 0.05 and ≥ 1-fold change, n = 1887) represent pulldown proteins enriched in neuron-TurboID cortex. Purple symbols (two-sided t-test, p ≤ 0.05 and ≥ 1-fold change, n = 679) represent pulldown proteins enriched in astrocyte-TurboID cortex. **j**. Transcriptome GO analysis of DEGs visualized in panel i. of overrepresented ontologies of neuron-TurboID (blue scale) and astrocyte-TurboID (purple scale) pulldowns, highlighting ontologies related to neuronal (e.g., neurotransmitter receptor activity and neuron to neuron synapse) and astrocyte function (e.g., lipid binding and astrocyte end-foot), respectively.

Following streptavidin pulldown, we determined protein enrichment using a silver stain (**Supplementary Fig. 4c and d**) and eluted high-quality RNA in both astrocyte-TurboID and neuron-TurboID cortical tissue, but negligible RNA in non-TurboID controls (**Fig. 2d**), confirming transcriptome enrichment only in the presence of biotinylation. We next prepared samples for LFQ-MS. Raw intensity and processed intensity values are found in Supplementary Data 16-18, respectively. Notably, MDS analysis of the proteomes showed separation between neuronal and astrocytic proteomes (**Fig. 2e**). We also confirmed the cell type specificity of the proteomes including enrichment of canonical astrocytic proteins e.g., Hepacam, Glu, Aqp4, Plpp3, p ≤ 0.05, ≥ 1-fold change) and GO terms (e.g., astrocyte end-foot, glial cell projection and astrocyte differentiation) in astrocytic pulldowns and neuronal proteins (e.g., Map2, Ncam1, Mapt, p ≤ 0.05, ≥ 1-fold change) and GO terms (e.g., synaptic signaling dendrite and neuron projection,) in neuronal pulldowns (**Fig. 2f and g**, Supplementary Data 19 and 20). These data are consistent with our prior work using astrocyte-TurboID and neuron-TurboID systems^34^. It is important to highlight that the lysis buffer used in the current studies was a homogenization buffer intended to maintain RNA-protein interactions, as opposed to 8 M urea lysis buffer in prior work^34^.

Like the proteome, we conducted MDS, differential enrichment and GO analysis of the neuronal and astrocytic transcriptomes. Raw RNA-seq and processed counts values are found in Supplementary Data 21 and 22, respectively. Like the proteome, MDS analysis identified distinct neuronal and astrocytic transcriptomes (**Fig. 2h**), each of which were enriched for canonical cell type specific markers (**Fig. 2i and j**, Supplementary Data 23 and 24), suggesting that the pulldowns are cell type enriched. To further validate the cell type specificity of the pulldown transcriptomes, we compared the relative gene expression of top astrocytic and neuronal genes from Zhang et al., 2014^51^ between the bulk cortex and the cell type enriched pulldowns (**Supplementary Fig. 5)**. The cell type enriched gene lists can be found in Supplementary Data 25. The astrocyte-TurboID pulldown transcriptomes exhibited a high expression of astrocytic genes (e.g., *Tnc, Sox9* and *Aqp4*) and a low expression of neuronal genes (e.g., *Tmem130, Slc10a4* and *Npy*) (**Supplementary Fig. 5a**), and the converse was true for the neuron-TurboID pulldown transcriptomes (**Supplementary Fig. 5b**).

We included a 5-day repeated LPS paradigm in our original astrocyte SPARO study to examine effects of LPS-induced neuroinflammation of astrocyte profiles (**Supplementary Fig. 6**). We observed modest effects of LPS at both the proteome (p ≤ 0.05, ≥ 1-fold change, n = 11 up with LPS, n = 35 up in non-LPS controls, Supplemental Data 18) and transcriptome levels (p ≤ 0.05, ≥ 1-fold change, n = 141 up with LPS, n = 41 up in non-LPS controls, Supplemental Data 21), suggesting that the LPS effect may have been partially washed out during the one-week interval between the last LPS dose and euthanasia (**Supplementary Fig. 6b and c**). Despite this, we observed modest yet meaningful effects of LPS on the astrocyte pulldown transcriptome using the SPARO approach, including upregulation of canonical reactive genes (e.g., *Lcn2* and *Serpina3g*, **Supplementary Fig. 6b**). We also performed GO analysis of DEGs and found synapse– and translation-related terms enriched with LPS treatment (**Supplementary Fig. 6d**, Supplementary Data 26). In contrast, LPS-induced changes were minimal at the level of the astrocyte proteome (**Supplementary Fig. 6c**), suggesting a discordance between LPS-induced effects on astrocytes at the transcriptomic and proteomic levels. We did not find any GO terms enriched from DEPs, due to the small number of DEPs found within each sample group.

These results demonstrate the application of the SPARO approach to obtain biologically meaningful proteomes and transcriptomes of astrocytes and neurons while retaining their native states in adult mouse cortex.

### The SPARO– and RiboTag-enriched astrocytic transcriptomes are similar

We next benchmarked SPARO against RiboTag, a gold-standard in vivo cell type-specific transcriptomic profiling method^38,39^. RiboTag mice express a hemagglutinin tag (HA) modified allele of the *Rpl22* gene (*Rpl22-HA*), a major component of the polyribosome complex^38,39^. In the presence of Cre recombinase, HA-tagged polyribosomes can be isolated from target cell types using an anti-HA antibody and immunoprecipitation methods. The mRNAs interacting with the ribosomes during translation can then be extracted and quantified to obtain a snapshot of actively translated transcripts (translatome) in specific cell types. To assess the similarity between the TurboID-enriched and RiboTag-enriched transcriptomes, we compared both methods using cortical Aldh1l1-expressing astrocytes in vivo (**Fig. 3a**). We bred Aldh1l1^CreERT2/wt^ mice with Rpl22^tm1.1Psam^ or TurboID mice to generate astrocyte-RiboTag and astrocyte-TurboID mice, respectively. We administered tamoxifen to astrocyte-TurboID, astrocyte-RiboTag and Aldh1l1^CreERT2/wt^ or Rpl22^tm1.1Psam^ (control) mice at 7 weeks of age, followed by a 3-week gap, then performed cortical tissue lysis, pulldown of biotinylated or HA-tagged proteins, and RNA-seq (**Fig. 3a**).

**Figure 3:**
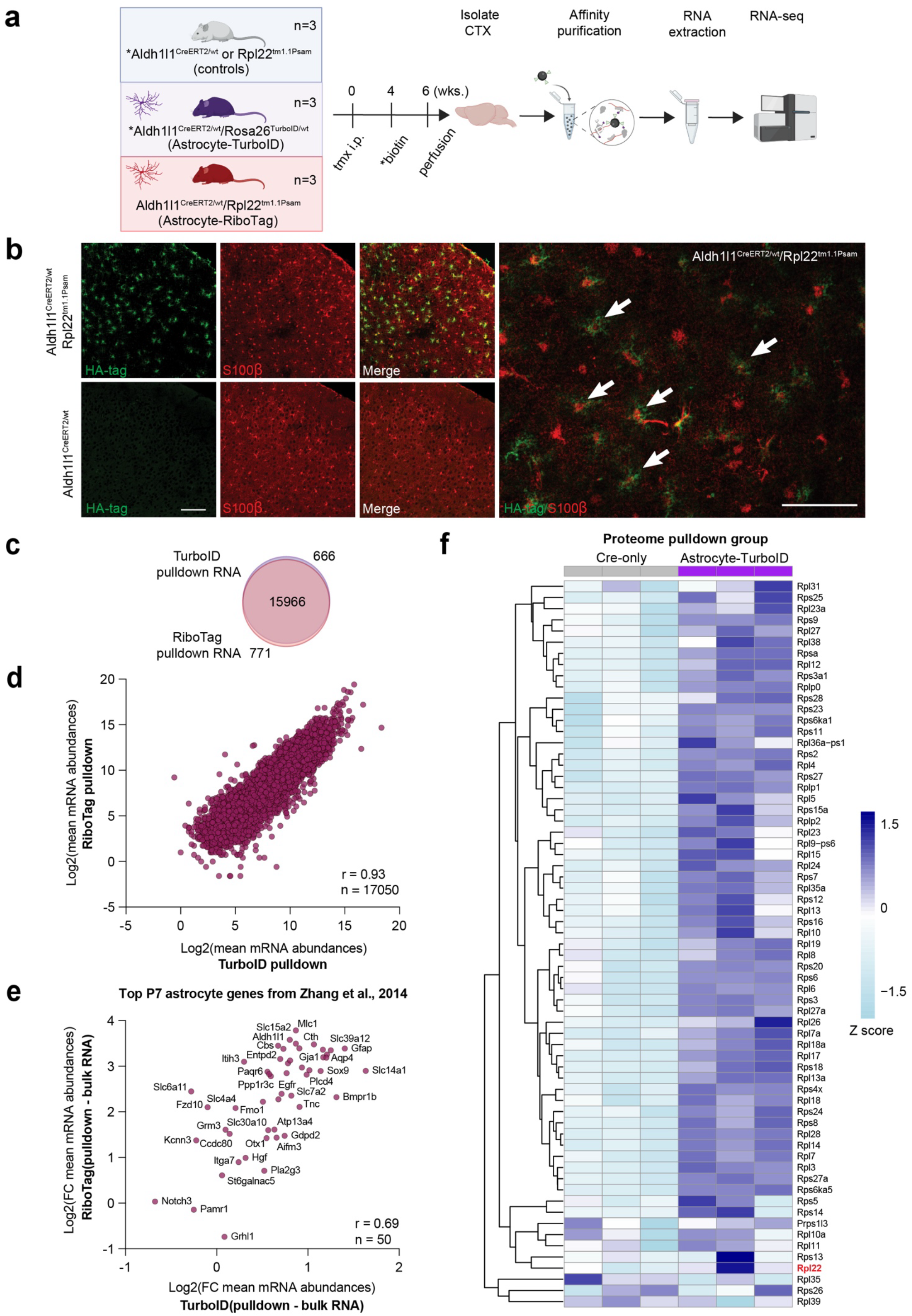
The TurboID-enriched transcriptome is comparable to the RiboTag-enriched transcriptome of cortical astrocytes. **a**. Experimental overview to compare the TurboID-enriched astrocyte transcriptome to the RiboTag-enriched astrocyte transcriptome. Aldh1l1^CreERT2/wt^ or Rpl22^tm1.1Psam^ (control, n = 3 mice), Aldh1l1^CreERT2/wt^/Rosa26^TurboID/wt^ (astrocyte-TurboID, n = 3 mice) and Aldh1l1^CreERT2/wt^/ Rpl22^tm1.1Psam^ (astrocyte-RiboTag, n = 3 mice) mouse models (aged 2.5-3.5 months). Astrocyte-TurboID, astrocyte-RiboTag and control mice received tamoxifen (75 mg/kg/day via intraperitoneal (i.p.) injections x 5 days). After 3 weeks astrocyte-TurboID mice received biotin-containing water (37.5 mg/L) for two weeks. Cortical tissue was isolated and lysed for affinity purification of either biotinylated ribosomal and RNA-binding proteins from astrocyte-TurboID mice, or HA-tagged ribosomal protein Rpl22, followed by RNA extraction and RNA-sequencing. **b**. Representative immunofluorescence images of Aldh1l1^CreERT2/wt^ control and astrocyte-RiboTag mice sagittal brain sections (n = 2-3 mice/experimental group) confirms the presence of the HA recombinant protein (green) in the astrocyte-RiboTag mouse cortex and not the controls. The HA tag colocalizes with the astrocytic marker, S100β (red), confirming astrocytic specificity. Scalebar = 50 μm. **c**. Venn diagram representation of mRNA numbers identified by RNA-seq present in the TurboID-enriched and RiboTag-enriched astrocyte pulldown transcriptomes. **d**. Scatter plot visualization comparing the expression of 17,050 transcripts present in the TurboID (x-axis) or RiboTag (y-axis) pulldown transcriptomes (Pearson correlation coefficient r = 0.93). Genes that are not present in both lists were removed from the analysis. **e**. Scatter plot visualization comparing the expression of the top 50 postnatal day 7 (P7) astrocyte-specific transcripts enriched in the TurboID (x-axis) or RiboTag (y-axis) pulldown transcriptomes after bulk RNA subtraction (Pearson correlation coefficient r = 0.69). The astrocyte-specific gene lists were acquired from Zhang et al., 2014. **f**. Heatmap visualization displaying the relative abundance (via Row Z score, purple = relative high abundance or enrichment and blue = relative low abundance or depletion) of ribosomal proteins (n = 65) present in the Aldh1l1^CreERT2/wt^ (Cre only) and astrocyte-TurboID pulldowns.

Immunofluorescent imaging of the cortex showed the presence of HA recombinant protein in astrocyte-RiboTag brains, which co-localized with the astrocytic marker, S100β (**Fig. 3b**). RNA extractions were of similarly high quality across all SPARO and RiboTag pulldowns, with low levels seen in the RiboTag only controls (**Supplementary Fig. 7a and b**). We next prepared samples for RNA-seq. Raw RNA-seq and processed counts values are found in Supplementary Data 27 and 28, respectively. Importantly, transcript overlap between the astrocyte-TurboID and astrocyte-RiboTag pulldowns was 95% (**Fig. 3c**). The cell type specificity of RiboTag and SPARO transcriptomes (**Supplementary Fig. 7c**), as well as the correlation across datasets (Pearson correlation coefficient r = 0.93, n = 17050 shared transcripts, **Fig. 3d**) were both robust. Next, we examined whether highly cell type-specific astrocytic genes from Zhang et al., 2014 were correlated across the TurboID– and RiboTag-enriched transcriptomes. We found that the mRNA abundances were highly correlated (Pearson correlation coefficient r = 0.69, n = 50 genes, **Fig. 3e**).

We also performed differential enrichment analysis and GO analysis between the astrocyte-TurboID and astrocyte-RiboTag pulldown transcriptomes (**Supplementary Fig. 7d-f**, Supplementary Data 29 and 30) and identified several differences in signatures. We hypothesize that these differences arise from the fact that RiboTag enriches all RNAs bound to ribosomal complexes that contain the ribosomal protein, Rpl22, whereas the SPARO approach captures mRNAs interacting with many biotinylated proteins that interact with RNA in astrocytes, which include 66 ribosomal proteins (including Rpl22) (**Fig. 3f**) and 67 RBPs (Supplementary Data 31).

### Correlation of the SPARO transcriptomes and proteomes of cortical astrocytes and neurons in vivo

We next evaluated the concordance between the in vivo proteomes and transcriptomes of astrocytes and neurons obtained using SPARO (**Fig. 4a**). The raw correlation between 1,934 mRNA and protein pairs was modest in both astrocytes (Pearson correlation coefficient r = 0.27, **Supplementary Fig. 8a**, Supplementary Data 32) and neurons (Pearson correlation coefficient r = 0.26, **Supplementary Fig. 8b**, Supplementary Data 33). To determine if these generally low correlations were simply a product of the complexity of an in vivo brain environment, we also performed similar transcriptome and proteome correlation analysis of the in vitro BV2-TurboID cells and found similar results (Pearson correlation coefficient r = 0.24, **Supplementary Fig. 8c**, Supplementary Data 34). We did observe the highest concordance between bulk cortical transcriptomes and proteomes (Pearson correlation coefficient r = 0.47, **Supplementary Fig. 8d**, Supplementary Data 35), which lack any cell type specificity.

**Figure 4:**
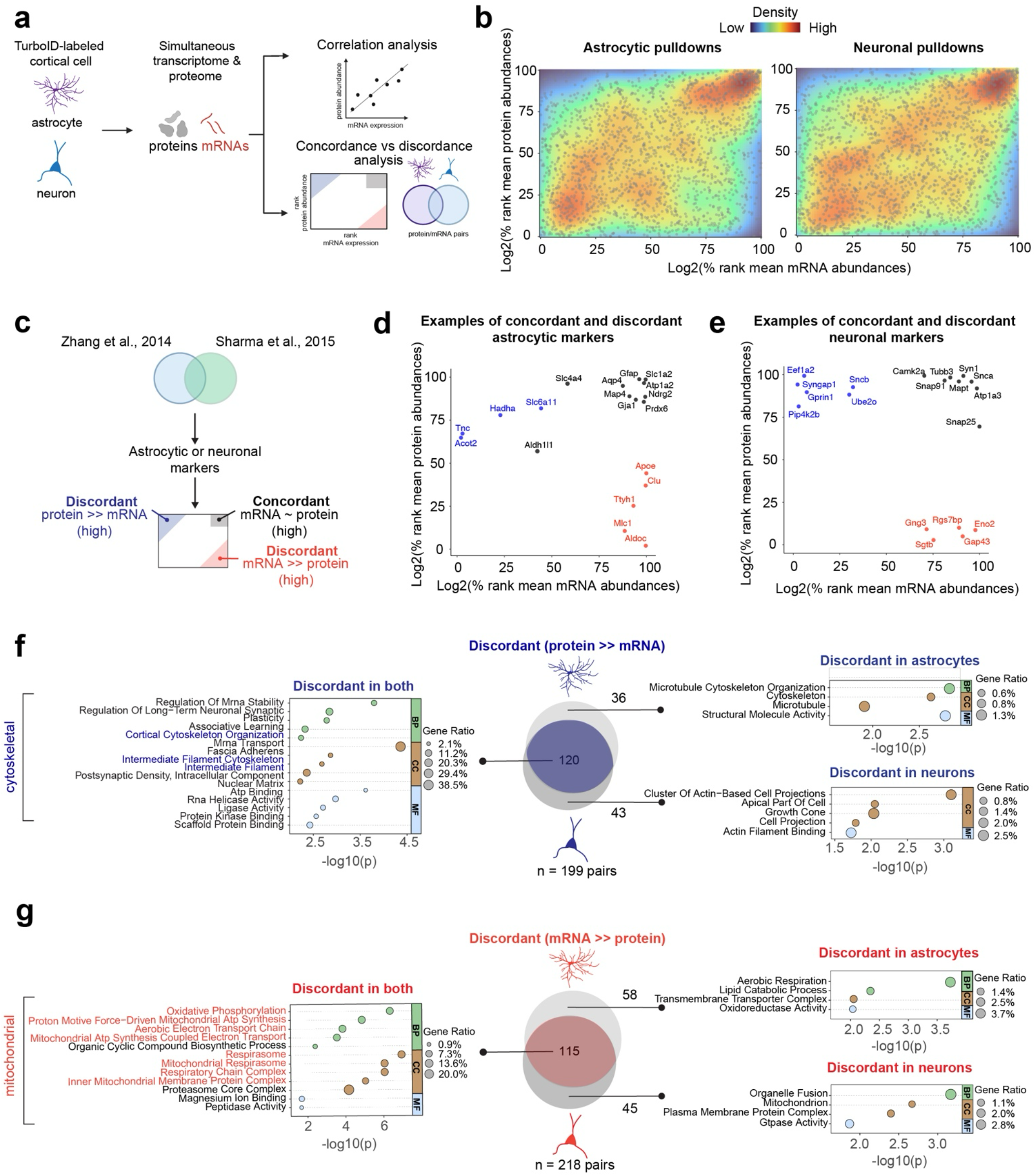
Correlation analysis of TurboID-based transcriptomes and proteomes of Aldh1l1-expressing astrocytes and Camk2a-expressing neurons in vivo. **a**. Schematic of analysis approach to compare SPARO of cortical astrocytes and neurons (n = 3 animals/group). **b**. Percent rank transformation and density plot visualization of 1934 gene and protein pairs between the astrocyte-TurboID (left) and neuron-TurboID (right) pulldown transcriptome (x-axis, via normalized, log_2_-transformed and percent ranked RNA-seq mean count values) and proteome (y-axis, via normalized, log_2_-transformed and percent ranked LFQ-MS mean intensity values). **c**. Schematic of analysis approach to nominate astrocytic and neuronal markers into highly concordant (mRNA ∼ protein in black) or highly discordant (protein >> mRNA in blue or mRNA >> protein in red) groups. The top astrocyte-specific and neuronal-specific markers were acquired from the unionization of the Zhang et al., 2014 and Sharma et al., 2015 datasets. **d**. Scatter plot visualization of examples of astrocytic markers that are concordant where mRNA and protein abundances are high and similar (top right, black, e.g., Gfap, Aqp4, and Slc1a2), discordant protein >> mRNA (top left, blue, e.g., Tnc, Acot2 and Hadha) and discordant mRNA >> protein (bottom right, red, e.g., Apoe, Tthy1 and Aldoc). **e**. Scatter plot visualization of examples of neuronal markers that are concordant where mRNA and protein abundances are similar (top right, black, e.g., Camk2a, Syn1, and Snca), discordant protein >> mRNA (top left, blue, e.g., Eef1a2, Syngap1, and Sncb) and discordant mRNA >> protein (bottom right, red, e.g., Gng3, Eno2, and Gap43). **f**. Venn diagram representation of mRNA and protein pair quantities present in the discordant (top left, blue, protein >> mRNA) group between astrocyte-TurboID and neuron-TurboID paired pulldown transcriptomes and proteomes. Gene-set enrichment analysis visualization of GO (via –log10(p value) of unique mRNA and protein pairs enriched in neurons and astrocytes. **g**. Venn diagram representation of mRNA and protein pair quantities present in the discordant (bottom right, red, mRNA >> protein) group between Aldh1l1-expressing astrocytic and Camk2a-expressing neuronal paired pulldown transcriptomes and proteomes. Gene-set enrichment analysis visualization of GO (via –log10(p value) of unique mRNA and protein pairs enriched in neurons and astrocytes.

Quantifying concordance is further complicated by innate technical differences between LFQ-MS and RNA-sequencing. For example, the dynamic range and sensitivity of both approaches differ vastly. Therefore, to better normalize against these biases, we calculated percentile ranked abundances for proteins and transcripts in each dataset (1,934 gene and protein pairs that overlap) and classified these mRNA-protein pairs into distinct groups based on level of mRNA-protein rank discordance (**Fig. 4b and Supplementary Fig. 8e**, Supplementary Data 32 and 33). In both astrocytes and neurons, and in the bulk cortex, we observed that most mRNA-protein pairs fell into either concordantly low or high groups (Log_2_ % rank values between 75 and 100 for both the x and y-axis). Together, these findings verify modest levels of concordance between protein and mRNA abundances^52–54^, although mRNAs that tend to be highly expressed in neurons and astrocytes, and in the cortex in general, also tend to be highly abundant at the proteomic level.

We next classified mRNA and protein pairs from the astrocyte and neuronal pulldowns into either concordant (mRNA ∼ protein, > 0.75 in black) or discordant (protein >> mRNA, > – 0.50 in blue, or mRNA << protein levels, > 0.50 in red) groups based on a rank differential of abundance values **(Supplementary Fig. 8f**, Supplementary Data 32 and 33). We wondered whether there were examples of canonical cell type-specific markers that fall into either highly concordant or discordant quadrants. To designate astrocytic and neuronal enriched canonical markers, we used a unionized marker list^55–57^ generated from the Zhang et al., 2014^51^ transcriptomic data and the Sharma et al., 2015^54^ proteomic data from acutely isolated astrocytes and neurons from the mouse brain (**Fig. 4c and Supplementary Fig. 8g and h**, Supplementary Data 36). A subset of example markers belonging to each category are shown in **Fig. 4d**. We repeated the same analysis for the neuronal pulldown proteomes and transcriptomes (**Fig. 4e**). These findings nominate cell type-specific astrocytic and neuronal markers that may be preferentially suitable for either transcriptomic or protein-based studies, as well as those that are highly abundant at both mRNA and protein levels.

mRNA-protein discordance can arise from many biological phenomena, which may potentially also vary across cell types. We next were curious wondered whether discordant proteins and mRNAs were consistent across astrocytes and neurons. We assessed the level of overlap in mRNA-protein pairs across astrocytes and neurons in 3 quadrants (high mRNA and high protein, high mRNA and low protein, low mRNA and high protein, Supplementary Data 37). Across all 3 groups, most concordant mRNA-protein pairs (high RNA, high protein) were present in both cell types (**Supplementary Fig. 8i**). Many discordant mRNA-protein pairs were also shared across cell types (**Fig. 4f and g**). Interestingly, we observed a pattern in which the types/categories of discordant genes were generally conserved across cell types, with specific subtypes of genes that are cell type specific. For example, within the discordant group where protein exceeded mRNA (protein >> mRNA) abundance, we found overrepresented ontologies of cytoskeletal pathways in both astrocytes and neurons. However, within astrocytes, only terms related to microtubules were enriched, as opposed to actin machinery in neurons (**Fig. 4f**). For the discordant group of genes in which mRNA exceeded protein (mRNA >> protein) abundance, we found a consistent signature in both neurons and astrocytes of ontologies related to mitochondria. However, within astrocytes this signature comprised of genes involved in aerobic respiration, whereas neurons were biased towards genes involved with mitochondrial fusion and GTPase activity (**Fig. 4g**). This pattern most likely reflects the metabolic differences that exist between these two cell types. After performing GO analyses on the mRNA and protein pairs in each concordant and discordant quadrant, we also looked at the overlap of the GO terms across the two cell types. The cell type-specific GO enrichment patterns of mRNA and protein pairs in the concordant and discordant groups generally matched our results from GO analyses at the level of mRNA-protein pairs (**Fig 4f and g, Supplementary Fig. 9**, Supplementary Data 38). After evaluating the correlation between the paired proteomes and transcriptomes of each TurboID model, we aimed to uncover new mechanisms mediating concordance and discordance between mRNA and protein abundances in the bulk cortex, astrocytes and neurons. We found no relationship between the level of mRNA-protein discordance with the average RNA traits: % GC content, number of exons and transcript length or protein half-life^58^ (**Supplementary Fig. 10**, Supplementary Data 39 and 40), suggesting that these mRNA characteristics and protein synthesis dynamics are unlikely on their own to explain the observed discordance.

Together, these findings emphasize an important observation about discordant mRNAs and proteins across these two cell types. In general, the broad classes of mRNAs and proteins that are highly discordant in abundance are well conserved between neurons and astrocytes. It is only in the nuanced subtypes of these gene classes where each cell type exhibits a uniquely discordant repertoire of proteins and transcripts.

## Discussion

Measuring transcriptomic and proteomic levels is important to identify cellular and molecular mechanisms of biology. Less than 5% of gene products exhibit cell type-specific expression patterns (via > 4-fold enrichment), and as a result, bulk tissue –omics studies cannot completely resolve changes occurring at the level of individual cell types^51,54^. Since proteome and transcriptome level abundances only modestly correlate with each other^52–54^, complementary profiling of both levels of abundances are needed. We describe a new approach to capture the cell type-specific transcriptome and proteome simultaneously that is applicable to in vitro and in vivo models. This method takes advantage of the biotin ligase, TurboID, to biotinylate cytosolic proteins that interact with RNA including ribosomal and RNA-binding proteins, which allows for enrichment of biotinylated proteins for proteomics as well as protein-associated mRNA for transcriptomics. This approach, called SPARO, addresses a major methodological gap in the field by providing a single pipeline for dual transcriptomics and proteomics from a desired cell type, while retaining the cell’s native state in the tissue. In this study, we establish the SPARO approach to obtain valid proteomes and transcriptomes from BV2 microglia in vitro and further capture the effect of a neuroinflammatory stimulus on microglia, using whole-cell bulk profiles of BV2 cells as gold standard references. Based on these in vitro results, we extended SPARO to the in vivo setting, using Camk2a-expressing neurons and Aldh1l1-expressing astrocytes as brain cell types of interest, further validating the method for in vivo applications. By comparing SPARO-derived with RiboTag-derived astrocyte transcriptomes, we further confirmed that both approaches yield comparable results. Leveraging simultaneously enriched transcriptomes and proteomes of neurons and astrocytes, we also investigated patterns of mRNA-protein concordance across cell types.

### Feasibility and validity of the SPARO approach

To first validate SPARO in vitro, we used a validated BV2-TurboID microglial cell line^35,49^ where we could benchmark our findings against the “ground truth” of a homogenous cell population. Consistent with previous work^35^, the TurboID-enriched pulldown proteome captured approximately 50% of proteins identified by bulk proteomics. The level of correlation between the bulk proteome and the cytosolic biotinylated proteome from BV2 microglia was also modest. We attribute these to the cytosolic bias of TurboID-NES used in our experiments, which under-samples nuclear and mitochondrial proteins^26^. More sensitive MS approaches are also likely to increase the depth of the proteome. The transcriptome of enriched mRNAs after streptavidin pulldown showed very high overlap and correlation with the bulk transcriptome, highlighting that the SPARO transcriptome is highly representative of the bulk transcriptome. As predicted, we found that mitochondrial DNA-encoded genes (e.g., *COX1, CYTB* and *ND2*) were under sampled in the pulldown transcriptomes, consistent with the inability of cytosolic TurboID-NES to label the intramitochondrial matrix compartment.

Using LPS as a neuroinflammatory stimulus to activate BV2 microglia, we used SPARO and verified LPS effect at the pulldown transcriptome level strongly reflected LPS effect at the bulk transcriptome level. At the proteomic level, LPS effect at the bulk level was modestly recapitulated by the pulldown level, potentially explained by the cytosolic bias of TurboID-NES. These observations highlight major differences in LPS effects at the proteomic and transcriptomic levels in BV2-microglia, although canonical markers of microglial activation induced by LPS tend to be conserved at both the protein and mRNA levels. Based on successful demonstration of the feasibility and validity of the SPARO approach in vitro in a mammalian cell system, we next applied the SPARO approach in vivo to contrast the native-state proteome and transcriptome of astrocytes and neurons from the adult mouse cortex. As seen in our in vitro studies, RNA bound to biotinylated proteins were enriched from astrocyte-TurboID and neuron-TurboID cortical tissues. We confirmed that the astrocyte– and neuron-TurboID pulldown proteomes and transcriptomes were cell type enriched via high enrichment of astrocytic or neuronal markers (e.g., Hepacam and Aqp4 in astrocytes and Map2 and Tmem130 in neurons) and GO terms related to their unique cellular functions. One important observation that we noted was that the effect of LPS on astrocytes in vivo (as well as BV2 cells in vitro) was far more dramatic at the transcriptomic level. This is well reflected by the enrichment of the canonical inflammatory reactive marker, *Lcn2*, at the transcriptomic level only, providing one potential explanation for why protein-based readouts of reactivity are oftentimes more difficult to demonstrate. Overall, these observations provide further validation that SPARO in vivo can be used to quantify cell-type differences at the RNA and protein levels.

### The SPARO with RiboTag transcriptomes are similar and complementary

Our benchmarking of SPARO with RiboTag noted some interesting differences. A notable difference between the SPARO and RiboTag transcriptomes is that in SPARO, TurboID biotinylates 133 proteins that interact with RNA (e.g., ribosomal and RBPs), while the RiboTag approach labels one ribosomal protein, suggesting that the TurboID-mediated SPARO approach captures more than ribosome-bound transcripts. Additionally, TurboID biotinylates the Rpl22 protein, suggesting that the transcripts associated with Rpl22 can be captured similarly to the RiboTag approach. When we compared the two methods using cortical Aldh1l1-expressing astrocytes, we found that the transcriptomes are comparable when looking at the presence and abundance of transcripts. Furthermore, the abundance of canonical astrocytic enriched genes (e.g., *Gfap, Aqp4, Sox9* and *Aldh1l1*) are highly correlated. When we differentially compared the RiboTag and SPARO-enriched transcriptomes, we found overrepresented ontologies involved in translation-, splicing– and mitochondrial-related terms in the RiboTag pulldowns and synapse– and ion transport-related terms in the SPARO pulldowns. These differences may be attributed to SPARO capturing mRNA species beyond only those that are actively being translated. Notably, the highest DEP in our SPARO astrocytic pulldown proteome is Aqp4, which is a highly abundant water channel found in the astrocytic end-feet^59^. Perhaps, the differences highlight methodological differences between the two approaches. For example, we observed that biotinylation by TurboID in astrocytes preferentially labels astrocytic end-feet, which is where Aqp4 is mostly localized. Despite these differences, we show that the SPARO and RiboTag astrocytic transcriptomes are indeed comparable and serve as appropriate tools to measure astrocytic genes in adult cortical astrocytes in vivo.

### SPARO to assess the concordance between paired proteomes and transcriptomes

One of the major advantages of SPARO is the ability to assess concordance between proteomes and transcriptomes of the same cells. In our in vivo astrocyte and neuron datasets, we observed only modest concordance between the pulldown proteomes and transcriptomes, which is in line with prior studies^52–54^. There are many possible reasons as to why the concordance is modest. From a technical perspective, the sensitivity of MS detection limits likely contributes to challenges of accurately measuring protein abundances and therefore, the correlation with transcript abundances. Additionally, transcriptomic and proteomic datasets are usually analyzed in isolation with different data processing and statistical analysis approaches that may contribute to the poor correlation between mRNA and protein levels. There are also limited resources to determine which allele of a gene yields a particular protein isoform. Moreover, the dynamic range of abundance is much higher for proteins than for transcripts^1,2,13^.

From a biological standpoint, other possibilities for the mRNA-protein discordance are post-transcriptional regulatory mechanisms such as mRNA and protein stability, mRNA half-life transcription and translation rates and efficiency, post-translational modifications, and vesicle-bound secretory pathways. Because we did not find a pattern associated with protein half-life and mRNA-protein discordance, we predict that other mechanisms or a collection of many cellular and molecular processes contribute to the discordance between mRNA and protein levels. We also observed that the BV2-TurboID bulk and pulldown transcriptomes in vitro are highly correlated, suggesting that the TurboID-enriched transcriptome is generally representative of the bulk. Therefore, the modest in vivo concordance between the pulldown proteomes and transcriptomes may be because of the cytosolic localization of TurboID and the biotinylation of a subset of the proteome. We identified astrocytic and neuronal markers that are highly concordant or discordant. Some of the discordant markers are derived from secreted proteins (e.g., Apoe and Gap43). Although cytosolic TurboID has been found to biotinylate secreted proteins^29,31,34,35,49^, it is not well understood if TurboID is able to localize within extracellular vesicles (EV) found in the endolysosomal secretory pathway. However, there is evidence that proteins can be secreted without being vesicle-bound^60^, which would enable cytosolic TurboID to biotinylate such proteins. Future studies using different TurboID constructs localized to various subcellular compartments or throughout the whole cell, will allow for a more wholistic quantification of the concordance between paired proteomes and transcriptomes using SPARO. Future studies that will investigate post-transcriptional regulatory processes within astrocytes and neurons in their native environment will also likely shed light on the determinants of discordance between mRNA and protein levels.

We also found that many of the cellular processes involved in the concordant and discordant mRNA and protein pair groups are shared between astrocytes and neurons. For example, in the discordant group (protein >> mRNA) astrocytes are enriched in microtubule proteins and neurons are enriched in actin proteins. It is not fully clear whether the abundance of these cytoskeletal proteins differ across the two cell types in general, across subcellular compartments or across different brain regions. It has been well reported that actin filaments are an important part of the neuronal cytoskeleton and contribute to the structure of axons and dendrites^61^. In contrast to our study in the cortex, a cytosolic BioID2-based LFQ-MS proteomics study found actin-filament-based processes enriched in mouse striatal astrocytes^33^, suggesting that there may be brain region differences. Another study found that microtubules are present in perivascular astrocytic endfeet^62^. Perhaps the high enrichment of microtubule proteins in the SPARO astrocytic pulldowns is due to capturing more end-feet-associated proteins, as suggested by the high abundance of Aqp4 protein as well. In the discordant group (mRNA >> protein), different mitochondrial-related terms were enriched in astrocytes and neurons. The transcripts associated with mitochondrial function are nuclear-encoded, as opposed to mitochondrial DNA-encoded. Therefore, we predict that the mitochondrial transcripts are higher across both cell types because SPARO utilizes a cytosolic TurboID and under samples mitochondrial proteins. Interestingly, we found ganglioside-induced differentiation associated protein 1 (Gdap1), a neuronal mitochondrial outer membrane protein implicated in Charcot-Marie-Tooth disease^63–65^, to be highly enriched in the SPARO neuronal transcriptome, suggesting that SPARO may be a useful tool to study disease-associated genes and proteins. Although the cellular processes are similar across the two cell types, our findings suggest that astrocytes and neurons may use different molecular machinery. The differences are likely due to the unique functions astrocytes and neurons play in the brain^66^.

### Broader applications of SPARO

While our studies provide the foundation for the SPARO approach and demonstrate its validity in vivo, the approach offers several future potential applications and extensions with broad implications within and beyond the neuroscience field. One interesting finding is the quantity of DEGs found in the astrocytic and neuronal pulldown transcriptomes. Nearly 3 times more DEGs were found in astrocytes when compared to neurons. The difference between cell types may be attributed to differences in the heterogeneity of neuronal and astrocytic populations, or differences in the abundance of enriched biotinylated proteins that interact with RNA (e.g., ribosomal proteins). Additionally, because astrocytes are highly metabolically active^81^ and since many of the top astrocytic DEGs are involved in metabolism-related processes (e.g., lipid binding), the SPARO method may be more efficient in capturing transcripts related to metabolism that are more highly abundant in astrocytes. Further research will be needed to parse out cell-type differences in transcript abundances and the biases of the SPARO method. Additionally, with the rapid evolution of MS instrumentation and MS pipelines, it is anticipated that the SPARO approach can be extended to small brain regions and subcellular compartments with significantly deeper proteomic coverage in future studies^67,68^.

TurboID biotinylates RBPs, therefore, SPARO also has the potential to quantify RNA species other than mRNAs including miRNAs, snRNAs, and other lncRNAs. Recent work has highlighted that lncRNAs play a significant role in various cellular processes including transcription, translation, metabolism and signaling^70^. Interestingly, *Xist*, a lncRNA that is important for mammalian females to transcriptionally silence one of the pair of X chromosomes^71^, was highly enriched in the neuronal pulldown transcriptome in which 2/3 of the mice were female. The astrocytic samples were derived from male mice. This observation highlights that SPARO may be a useful tool to study sex differences at the transcriptomic and proteomic levels. Lastly, future modifications of TurboID in vivo models using both transgenic^72^ and AAV-based approaches^33,73^, may provide opportunities to restrict TurboID to other cellular compartments or remove the cytosolic restriction by excluding the NES tag. These future endeavors are likely to extend the SPARO approach to interrogate compartment-specific protein and mRNA processes such as local translation that occurs in synapses or glial processes^74–80^. TurboID-EV was recently developed^69^ and a TurboID-EV-SPARO approach will be an exciting future tool to better understand the RNA and protein compositions of EVs.

In conclusion, SPARO is an exciting new tool that can be used to study many facets of cellular and molecular biology. For example, SPARO in vitro can be used for cause-and-effect mechanistic studies using reductionist models in multi-cellular model systems. Additionally, SPARO can be used to study how transcriptional and translational processes are fully regulated, changes in the efficiency of protein biosynthesis, processes that contribute to the discordance between mRNAs and proteins, mRNA and protein degradation, and much more. A major advantage of SPARO is that it can be adapted to study different cell types, brain regions, and other tissues outside the brain, making the approach a highly versatile and broadly applicable tool for the field of molecular biology.

## Methods

### In vitro studies

Mouse microglia BV2 cells were cultured in 0.2 μm vacuum filter sterilized Dulbecco’s Modified Eagle Medium supplemented with high glucose and L-glutamine containing 1% penicillin-streptomycin, and 10% fetal bovine serum. Cells were incubated in 100 mm cell culture dish at 37°C and 5% CO_2_ until reaching 80-90% confluency. Cells were treated with 100 ng/mL of LPS for 48 h and 200 μM of biotin for 24 h during the second day of LPS treatment.

### In vivo studies

Mice were housed in the Division of Animal Resources vivarium at Emory University under a standard environment (12 h light/ 12 h dark cycle, temperature 72°F, humidity range 40-50%) with access to food and water ad libitum. All animal-related studies (PROTO-201700821) were conducted with the approval of Emory’s Institutional Animal Care and Use Committee following the National Institute of Health’s “Guide for the Care and Use of Laboratory Animals” and reported in accordance with the ARRIVE guidelines.

Transgenic approach for Cre recombinase expression in neurons and astrocytes using Rosa26^Tur-^ boID/wt mice:

### Neuronal TurboID cohort

Rosa26^TurboID/wt^ (Jackson Labs, Strain No. 037890) were crossed with Camk2a^Cre-ERT2^ (Jackson Labs, Strain No. 012362) mice to obtain heterozygous Rosa26^TurboID/wt/^Camk2a^CreERT2/wt^ (“neuron-TurboID”, n = 3, 1 M and 2 F littermate mice. Heterozygous littermate mice Camk2a^CreERT2/wt^ (“Camk2a”, n = 2, 1 M and 1 F) were used as controls.

### Astrocytic TurboID cohort

Rosa26^TurboID/wt^ (Jackson Labs, Strain No. 037890) were crossed with Aldh1l1^CreERT2^ (Jackson Labs, Strain No. 031008) mice to obtain heterozygous Rosa26^TurboID/wt^/Aldh1l1^CreERT2/wt^ (“astrocyte-TurboID”, n = 8, 6 M and 2 F). Heterozygous littermate mice Aldh1l1^CreERT2/wt^ (“Aldh1l1”, n= 3, 2 M and 1 F) were used as controls. For astrocyte-TurboID LPS studies, systemic LPS at 0.75 /kg/dose x 5 days was given via i.p. injection during the first week of biotin water supplementation. Based on previous work and our own, we validated neuron– and astrocyte-specificity and nonleaky Cre activity of the Camk2a^CreERT2^ and Aldh1l1^CreERT2^ models, respectively^40,50^.

### Astrocytic RiboTag cohort

RiboTag mice, Rpl22^tm1.1Psam^ (Jackson Labs, Strain No. 029977) were crossed with Aldh1l1^CreERT2^ (Jackson Labs, Strain No. 031008) mice to obtain heterozygous Rpl22^tm1.1Psam^/Aldh1l1^CreERT2/wt^ (“astrocyte--RiboTag”, n = 3, 3 F). Heterozygous littermate mice Rpl22^tm1.1Psam^ (“RiboTag”, n= 2, 1 M and 1 F) were used as controls.

All mice were given tamoxifen (75 mg/kg) intra-peritoneally (i.p.) for 5 days at 6 weeks of age and allowed 3 weeks of Cre recombination. After Cre recombination, TurboID mice were given water supplemented with biotin (37.5 mg/L) for 2 weeks until euthanasia at 3-4 months of age. RiboTag mice were euthanized at 2.5 months of age. Mice from each cohort were anesthetized using ketamine/xylazine (ketamine 87.5 mg/kg, xylazine 12.5 mg/kg) followed by transcardial perfusion with ice-cold 1X PBS. The brain was immediately removed and hemisected along the mid-sagittal line. The left hemisphere was immediately dissected to obtain the cortex and then snap frozen using dry ice. The right hemisphere was used for immunohistological studies.

### Immunohistochemistry

For immunohistochemistry studies, mice were euthanized and perfused with ice-cold 1X PBS, and the brain was dissected. One hemisphere was immediately transferred into 4% PFA (Cat No. J19943.K2) and fixed overnight for immunohistological analysis. After PFA fixation, the brain was washed with 1X PBS 3 times to remove residual PFA and transferred into 30% sucrose solution until sectioning. After removing extra sucrose, the fixed and sucrose-saturated hemisphere was embedded in the Tissue-Tek optimal cutting temperature compound (Cat No. 4583, Sakura) and frozen on dry ice. Serial 40 μm sagittal sections of the brain were generated using a cryostat (Leica Biosystems, CM1850) and stored free-floating in cryoprotective media (Glycerin, Ethylene Glycol and 0.1 M Phosphate Buffer in ratio of 2.5: 3.0: 5.0) at –20°C. For immunofluorescence (IF) staining, 3-4 brain sections from each mouse were washed 3 times with 1X TBS and then blocked and permeabilized by incubating in 1X TBS containing 0.25% Triton X-100 (TBS-T) and 5% goat serum for 1 h at room temperature. Following, sections were incubated with primary antibodies diluted in 1X TBS-T containing 1% goat serum overnight at 4°C. Next, the sections were washed 3X with 1X TBS and then incubated with fluorophore-conjugated secondary antibodies for 2 h at room temperature in the dark. All primary and secondary antibodies were used at dilution 1:500 including Streptavidin Protein, DyLight™ 488. Brain sections were then incubated for 5 minutes in nuclear staining reagent DAPI (1 μg/mL) dissolved in 1X TBS. The samples were further washed 3X with 1X TBS, mounted onto glass slides and coverslipped using ProLong Diamond Antifade Mountant (P36965, ThermoFisher). Images of the same brain regions for each mouse were captured using a Keyence fluorescence microscope (Keyence BZ-X810). Images were processed and analyzed using Image J software (FIJI Version 2.14).

### Cell and tissue homogenization

BV2 control and BV2-TurboID cells (1.0 x 10^6^ cells) were washed three times with cold 1X PBS containing 100 μg/mL of cycloheximide. Next, cells were incubated with 1 mL of cold homogenization buffer (10 mM HEPES pH 7.4, 150 mM KCl, 10 mM MgCl_2_ supplemented with 0.5 mM DTT, 0.1% v/v RNasin, 0.1% v/v SUPERasin, 1X Halt^TM^ protease and phosphatase inhibitors, 100 μg/mL cycloheximide) for 10 minutes, scaped and collected into a 1.5 mL Eppendorf LoBind tube. For brain samples, frozen Camk2a and Aldh1l1 controls and neuron-TurboID and astrocyte-TurboID mouse cortices (up to 50 mg) were lysed in same cold homogenization buffer as the BV2 cells with RNase-free glass beads and a Bullet Blender Homogenizer (Next Advance). RiboTag control and astrocyte-RiboTag mouse cortices were lysed in cold RiboTag homogenization buffer (50 mH Tris-HCl pH 7.5, 100 mM KCl, 12 mM MgCl_2_, 1% NP-40, 1 mM DTT, 1% v/v sodium deoxycholate 0.1% v/v RNasin, 0.1% v/v SUPERasin, 1X Halt protease and phosphatase inhibitors, 100 μg/mL cycloheximide) similarly to the TurboID samples. All homogenates underwent 3 rounds of sonication (5 s of active sonication at 25% amplitude followed by a 10 s incubation on ice) followed by centrifugation for 10 min at 2000 x g at 4°C. The supernatants were transferred to a new tube and protein concentrations were calculated using a Pierce BCA assay for proteomic studies.

### Immunoblotting

To confirm the presence of biotinylated proteins, 17 μg of cell or tissue lysates were resolved on a 4-12% Bis-Tris gel, transferred onto a nitrocellulose membrane, blocked with StartingBlock for 30 min and then probed with a streptavidin-AlexaFluor 680 antibody (ThermoFisher, S32358, dilution: 1:10,000 in StartingBlock) for 1 h at room temperature. To confirm TurboID construct presence, the membrane was probed with a rabbit anti-V5 primary antibody (dilution: 1:500 in StartingBlock) overnight and then incubated with a goat antirabbit HRP-conjugated secondary antibody (Jackson ImmunoResearch, 111-035-003, dilution: 1:10,000 in StartingBlock). Immunoblots were imaged with an Odyssey Infrared Imaging System (LI-COR Biosciences), or BioRad chemiluminescence system.

### Biotinylated protein and ribosomal protein enrichment for transcriptomics

Following confirmation of biotinylated proteins and V5 presence via immunoblotting, biotin-tagged proteins were enriched for using streptavidin-coated magnetic beads as described previously^34^. For each sample, 25 μL of streptavidin beads (ThermoFisher, 88817) in a 1.5 mL Eppendorf LoBind tube were washed 3 times with 1 mL of homogenization buffer on rotation for 2 min. The streptavidin beads were incubated with 300 μg of protein lysate from each sample with additional wash buffer (10 mM HEPES pH 7.4, 150 mM KCl, 10 mM MgCl_2_ supplemented with 0.5 mM DTT, 1% v/v NP-40 substitute, 0.1 % v/v RNasin, 0.1% v/v SUPERasin, 1X Halt^TM^ protease and phosphatase inhibitors, 100 μg/mL cycloheximide) to create a final volume of 250 μL on rotation for 1 h at 4°C. Next, the beads were washed four times with cold high salt buffer (10 mM HEPES pH 7.4, 350 mM KCl, 10 mM MgCl_2_ supplemented with 0.5 mM DTT, 1% v/v NP-40 substitute, 0.1 % v/v RNasin, 0.1% v/v SUPERasin, 1X Halt^TM^ protease and phosphatase inhibitors). After RiboTag control and Astrocyte-RiboTag cortical lysis, 800 μL of lysate was incubated with 4 μL of anti-hemagglutinin (HA) antibody (Biolegend, 901513) for 4 hours on end-over-end rotation at 4°C. For each sample, 25 μL of A/G beads (ThermoFisher, 88803) in a 1.5 mL Eppendorf LoBind tube were washed 3 times with 400 μL of RiboTag homogenization buffer. The A/G beads were incubated with 100 μL of HA-tagged lysate overnight on end-over-end rotation at 4°C. Next, the A/G beads were washed four times with cold RiboTag high salt buffer (50 mM Tris-HCl pH: 7.5, 300 mM KCl, 12 mM MgCl_2_, 1% v/v NP-40, 0.50 mM DTT, 100 μg/mL cycloheximide).10 mM HEPES pH 7.4, 350 mM KCl, 10 mM MgCl_2_ supplemented with 0.5 mM DTT, 1% v/v NP-40 substitute, 0.1 % v/v RNasin, 0.1% v/v SUPERasin, 1X Halt^TM^ protease and phosphatase inhibitors). After final washes, the streptavidin and A/G beads were resuspended in 100 μL of wash buffer and then added to 700 μL of Trizol and stored at –80°C until RNA extraction. For bulk/input lysate/animal, 50 μL of lysate remained at 4°C during the protein enrichment protocols.

### RNA extraction

After protein enrichment using streptavidin and A/B beads, RNA was extracted from the beads and or bulk lysates using a miRNeasy Mini Kit (Qiagen Inc., 217004) following the manufacturer’s instructions. RNA was eluted in RNase-free water and analyzed for purity and concentration using a Bioanalyzer (Agilent) prior to RNA-sequencing library preparation.

### RNA-sequencing and processing

After assessing RNA quality, cDNA library preparation was performed. The SMART-Seq^®^ v4 PLUS Kit (Takara Bio) was used following the manufacturer’s protocol. Bulk sequencing of 40 million paired-end reads per sample was completed using the Illumina-based platform at Admera Health. FASTQ files were evaluated for quality using FastQC^82^ and then mapped to the mouse genome, GRCm38 (mm10) using the Spliced Transcripts Alignment to a Reference (STAR)^83^ aligner with the paired-end option. The *featureCounts*^84^ package was used to generate a raw counts data matrix from mapped reads. Lowly abundant transcripts across all samples (average count value ≤ 10) were filtered out. After generating the filtered, raw counts data matrices, we used *DESeq2*^85^ to normalize the matrices and perform differential gene expression-based statistics.

### Biotinylated protein enrichment for proteomics

For each sample, 42 μL of streptavidin beads in a 1.5 mL Eppendorf LoBind tube were washed 2 times with 1 mL of RIPA lysis buffer (50 mM Tris, 150 mM NaCl, 0.1% SDS, 0.5% sodium deoxycholate, 1% Triton X-100) on rotation for 2 min. The beads were incubated with 500 μg of protein lysate from each sample with additional RIPA lysis buffer to create a final volume of 500 μL on rotation for 1 h at 4°C. Next, the beads were quickly centrifuged, placed on a magnetic rack and the supernatant was transferred to a new 1.5 mL Eppendorf LoBind tube and stored at –80°C. The beads were washed with the following buffers on rotation at room temperature: twice with 1 mL of RIPA lysis buffer for 8 min, once with 1 M KCl for 8 min, once with 1 mL 0.1 M sodium carbonate for ∼10 s, once with 1 mL 2 M urea in 10 mM Tris-HCl (pH 7.6) for ∼10 s, and twice with 1 mL RIPA lysis buffer for 8 min. After the final RIPA wash, the beads were resuspended in 1 mL of 1X PBS, transferred to a new tube and washed two more times with 1X PBS on rotation for 2 min. To confirm biotinylated protein enrichment, 10% of the streptavidin bead volume was transferred to a new 1.5 mL Eppendorf LoBind tube and boiled in 30 μL of 2X Laemmli protein loading buffer (Bio-Rad, 1610737) supplemented with 2 mM biotin and 20 mM dithiothreitol (DTT) at 95°C for 10 min to elute the biotinylated proteins. Following, 1/3 or the eluate was resolved on a 4-12% Bis-Tris gel, transferred onto a nitrocellulose membrane, blocked with StartingBlock for 30 min and incubated overnight with a rabbit anti-V5 antibody (Abcam, ab206566) on a shaker at 4°C. Next, the membrane was incubated with a goat anti-rabbit HRP-conjugated secondary antibody (Jackson ImmunoResearch, 111-035-003, dilution: 1:10,000 in StartingBlock) and streptavidin-AlexaFluor 680 antibody (ThermoFisher, S32358, dilution: 1:10,000 in StartingBlock) for 1 h at room temperature. The remaining 2/3 of protein eluate was resolved on a 4-12% BisTris gel for a silver stain (ThermoFisher, 24612). Immunoblots were imaged with an Odyssey Infrared Imaging System (LI-COR Biosciences), or BioRad chemiluminescence system. Silver-stained gels were imaged using a Canon scanner (CanoScan LIDE 300).

### Protein digestion

To prepare enriched biotinylated proteins for MS-based proteomics, the remaining 90% volume of the streptavidin beads were resuspended in 50 mM ammonium bicarbonate (Na_3_CO_2_). Next, the bound proteins were reduced with 1 mM DTT for 30 min at room temperature and then alkylated with 5 mM iodoacetamide (IAA) in the dark for 30 min. Following, proteins were subsequently digested overnight with 0.5 μg of lysyl (Lys-C) endopeptidase (Wako, 127-06621) and then 1 μg of trypsin (ThermoFisher, 90058) on the shaker at room temperature. After digestion, the resulting peptide mixtures were acidified to a final concentration of 1% formic acid and 0.1% trifluoroacetic acid, desalted with an HLB column (Waters, 186003908), and dehydrated using a vacuum centrifuge (SpeedVac Vacuum Concentrator). To prepare bulk lysates for MS-proteomics, 50 μg of protein from each sample was reduced in 5 mM DTT for 30 min at room temperature and then alkylated with 10 mM IAA for 30 min in the dark. After, each sample was diluted (4-fold) with 50 mM ammonium bicarbonate (ABC) buffer and then digested with 1 μg of Lys-C endopeptidase on the shaker overnight at room temperature. Next, samples were diluted (4-fold) with 50 mM ABC buffer and digested with 2 μg of trypsin on the shaker overnight at room temperature. After digestion, the peptide mixtures were acidified, desalted, and dried down as described above.

### Mass spectrometry

Dried peptides were prepared for liquid chromatography as described previously^34^. Following, an Orbitrap Lumos Tribrid MS with a high-field asymmetric waveform ion mobility spectrometry (FAIMS Pro)^86^ interface was used to obtain all mass spectra at a compensation voltage of –45V with similar parameters as done before^34^.

### Protein identification and quantification

Raw MS files for bulk and pulldown protein from BV2-TurboID in vitro and astrocyte– and neuron-TurboID in vivo studies along with respective controls were searched separately using the Andromeda^87^ search engine integrated into MaxQuant^88^ (version v2.4.2.0), against the 2020 mouse UniProt database (https://www.uniprot.org/help/reference_proteome) including sequences for V5 and TurboID. Search parameters including: variable and fixed modifications, allowed miscleavages, minimum/maximum peptide mass and length, mass tolerance for fragment and precursor ions and quantification settings were determined as done previously^34,35^. Peptide spectral match false discovery rate (FDR) was set to 1%. The MaxQuant output data (raw intensity values) were uploaded to R (version 4.3.1) for normalization, log_2_ transformation, and differential abundance analyses. To account for endogenously biotinylated and non-specific proteins captured during streptavidin affinity purification, mean intensity values from TurboID-negative control pulldown samples were subtracted from the TurboID-positive pulldown samples by group. To account for variability in TurboID expression, biotinylated protein abundance, and streptavidin affinity purification efficiency, intensity values were normalized depending on the dataset. For BV2 in vitro studies, abundance values from bulk protein and enriched pulldown samples were normalized by taking each sample’s log_2_-transformed summed protein intensity median and setting it to zero. For astrocyte-TurboID and neuron-TurboID in vivo studies, enriched pulldown samples were normalized by subtracting the difference from the mean of the log_2_ transformed summed protein intensity for 71 ribosomal subunit proteins present in all TurboID-positive pulldown samples. Proteins were filtered out if they had missing values in 2/3 samples across groups. Missing values were imputed from normal distribution for each sample. Nomination of proteins as either ribosomal or RNA-binding (via RNA-binding motif or RRM) were determined using the HUGO Gene Nomenclature Committee database (https://www.genenames.org/).

### Data analysis and visualization

We utilized differential enrichment analysis, multidimensional scaling (MDS), and gene set enrichment analysis (GSEA) to analyze transcriptomic and proteomic data from in vitro and in vivo studies. Data analysis and visualization was completed using R software (version 4.3.1) and Prism (GraphPad, version 10). For transcriptomic studies, the *DESeq2*^85^ Wald test was used to identify DEGs between groups. For proteomic studies, unpaired 2-tailed (equal variance assumption) T-test equivalent calculations using F value from ANOVA run strictly with 2 comparison groups were performed with the parANOVA suite of functions in R (https://github.com/edammer/parANOVA) to identify DEPs between groups. DEGs and DEPs are presented as volcano plots. GSEA was applied using the GOparallel function (https://github.com/edammer/GOparallel) that scrapes monthly updated .GMT formatted GO from the Bader Lab website (https://baderlab.org/) as gene symbol lists submitted to one-tailed Fisher’s exact test for GO enrichment of overlapping symbols in gene lists of interest, including DEPs and DEGs. GO terms meeting Fisher exact significance of p value ≤ 0.05 (i.e., a Z-> 1.96) were considered. Protein and gene input lists were significantly differentially biochemically enriched over bulk or background, in addition to gene lists that were different by treatment condition (p ≤ 0.05, ≥ 1 log_2_-fold change). To assess average GC content, number of exons, and average transcript length of transcriptomic data, we used the *biomaRt* R package^89,90^ and the BioMart/Ensembl databases^91^. To examine protein half-life information of proteomic data, we used values from Fornasiero et al., 2018^58^.

## Data availability

The MS proteomics data have been deposited to the ProteomeXchange Consortium^92^ via the PRIDE^93^ partner repository with the dataset identifier PXD059818. Transcriptomics data is available on the NIH Gene Expression Omnibus (GEO) repository with the dataset identifier GSE287770. Source data are provided with this paper.

## Acknowledgements

Research reported in this publication was supported by the National Institute on Aging and the National Institute of Mental Health of the National Institutes of Health: 1F31AG079597-01A1 (C.C.R.), R01AG075820 (N.T.S., S.R.), R01AG071587 (S.R.) and R01MH125956 (S.A.S.). Research was also supported in part by the Emory Integrated Proteomics Core (EIPC) which is subsidized by the Emory University School of Medicine and is one of the Emory Integrated Core Facilities. Additional support was provided by the Georgia Clinical & Translational Science Alliance of the National Institutes of Health under Award Number UL1TR002378 (EIPC). Additional support was provided by the National Center for Advancing Translational Sciences of the National Institutes of Health under Award Number UL1TR000454.

## Author contributions

Conceptualization: C.C.R., S.A.S., S.R., Methodology: C.C.R., E.B.D., H.X., L.C., P.K., C.E.G., M.M.S., R.S.N., S.M., D.K., R.K., Q.G., P.B., D.M.D., N.T.S., S.A.S., S.R., Investigation: C.C.R., E.B.D, N.T.S., S.A.S., S.R., Writing-Original draft: C.C.R., S.A.S., S.R, Writing-Review and Editing: C.C.R., E.B.D., H.X., L.C., P.K., C.E.G., M.M.S., R.S.N., S.M., D.K., R.K., Q.G., P.B., D.M.D., N.T.S., S.A.S., S.R., Funding acquisition: C.C.R., N.T.S., S.A.S., S.R., Resources: N.T.S., S.A.S., S.R., Supervision: S.A.S., S.R.

