## Supplementary Figures for "Simultaneous profiling of native-state proteomes and transcriptomes of neural cell types using proximity labeling"

Proximity labeling | transcriptomics | proteomics | microglia | astrocytes | neurons | mRNA-protein concordance

S.A.S

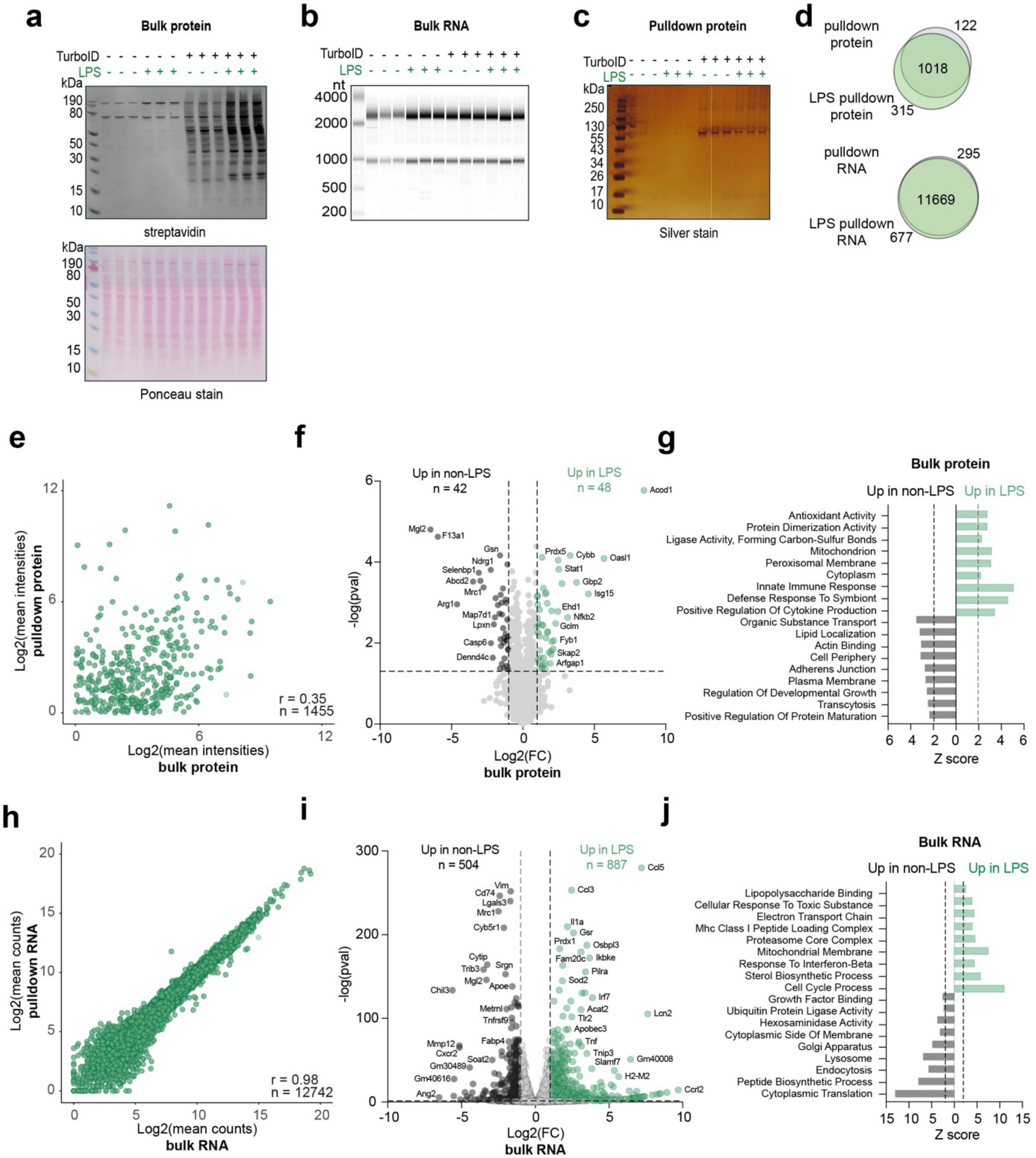


**Supplementary Figure 1: In vitro validation of SPARO in LPS-stimulated BV2-microglial cells.**
**a.** Immunoblot visualization (top) of biotinylated proteins from BV2-TurboID cells at the bulk level not seen in non-TurboID control cells (probed with streptavidin-680). Ponceau stain visualization (bottom) of protein abundance of BV2 control and BV2-TurboID bulk lysates showing equal protein loading for immunoblot analysis. **b.** RNA gel electrophoresis of RNA from BV2-TurboID bulk samples. High levels of rRNA and mRNA were detected in all BV2 samples. **c.** Silver stain visualization of enriched biotinylated proteins after affinity purification from a 10% aliquot of the streptavidin beads. Minimal protein enrichment was found in the BV2 control samples, and several proteins were enriched in the BV2-TurboID samples. **d.** Venn diagram representation of protein numbers identified by LFQ-MS present in the non-LPS-treated and LPS-treated BV2-TurboID pulldown proteomes (top). **e.** Correlation analysis of protein abundance from normalized LFQ-MS mean intensity values between the BV2-TurboID bulk and pulldown proteomes (n =

1455 proteins; Pearson correlation coefficient r = 0.35). **f.** Volcano plot representation of differentially enriched proteins (DEPs) between non-LPS and LPS-treated BV2-TurboID cells at the bulk level (n=3/group). Black symbols (two-sided t-test; $p \leq 0.05$ and ≥ 1-fold change; n = 42) represent DEPs in non-LPS-treated BV2-TurboID cells. Black symbols (two-sided t-test; $p \leq 0.05$ and ≥ 1-fold change; n = 48) represent DEPs in LPS-treated BV2-TurboID cells. **g.** Proteome GO analysis of DEPs visualized in panel f. of overrepresented ontologies of non-LPS treated (black) and LPS-treated (green) BV2-TurboID cells at the bulk level. Immune-related terms (e.g., innate immune response and positive regulation of cytokine production) are enriched in the LPS-treated samples. **h.** Correlation analysis of RNA abundances from normalized RNA-seq mean count values between the BV2-TurboID bulk and pulldown transcriptomes (n = 12742 genes; Pearson correlation coefficient r = 0.98). **i.** Volcano plot representation of differentially expressed genes (DEGs) between non-LPS and LPS-treated BV2-TurboID cells at the bulk level (n=3/group). Black symbols (two-sided t-test; $p \leq 0.05$ and ≥ 1-fold change; n = 504) represent DEGs enriched in non-LPS-treated BV2-TurboID cells. Green symbols (two-sided t-test; $p \leq 0.05$ and ≥ 1-fold change; n = 887) represent DEGs enriched in LPS-treated BV2-TurboID cells. **j.** Transcriptome GO analysis of DEGs visualized in panel f. of overrepresented ontologies of non-LPS treated black) and LPS-treated BV2-TurboID cells (green scale) at the bulk level. Immune-related terms (e.g., lipopolysaccharide binding and response to interferon-beta) are enriched in the LPS-treated samples.

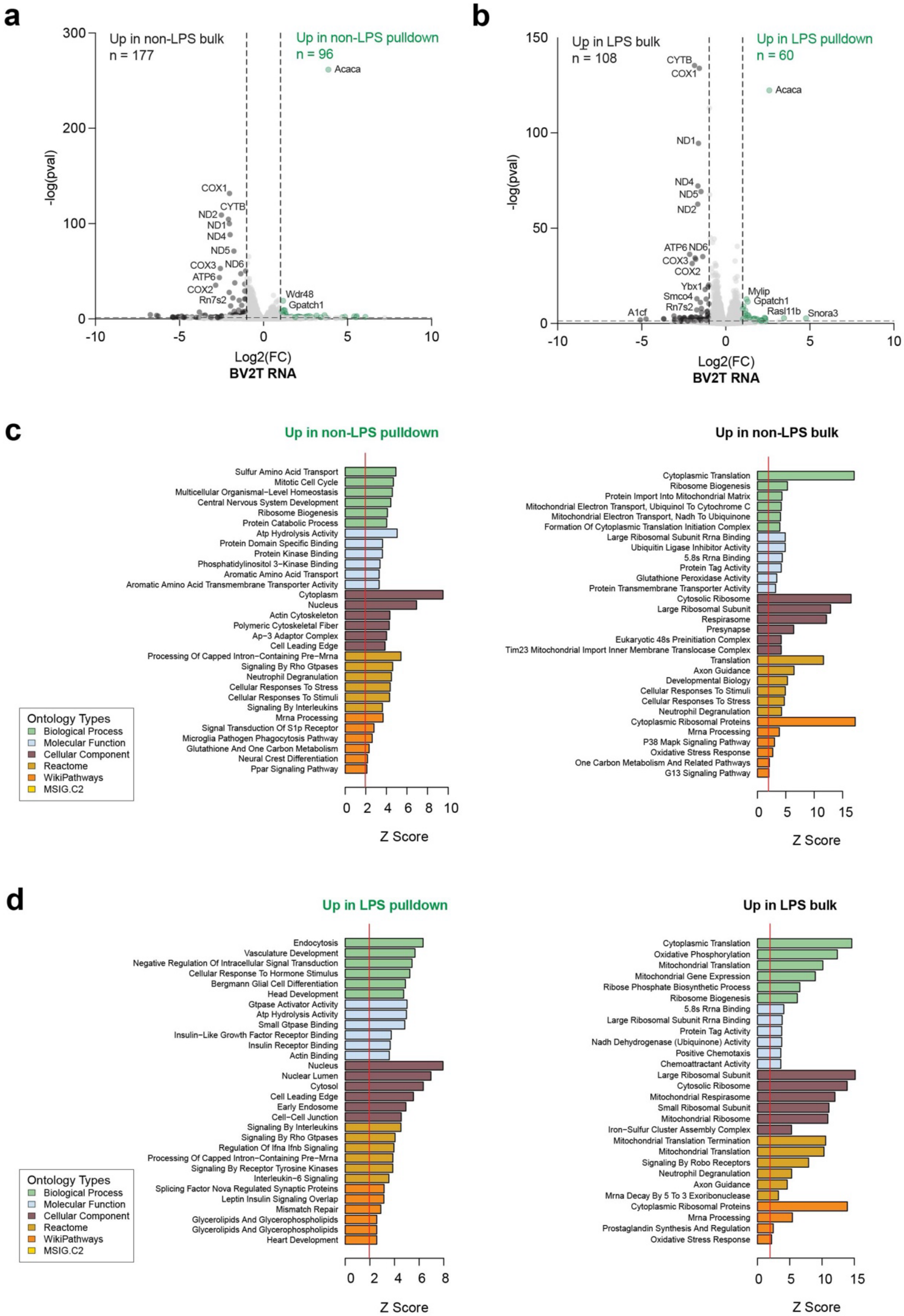


**Supplementary Figure 2: In vitro validation of the SPARO-enriched transcriptome in untreated and LPS-stimulated BV2-microglial cells.**

**a.** Volcano plot representation of DEGs between non-LPS treated BV2-TurboID cells at the pulldown and bulk level (n=3/group). Green symbols (two-sided t-test; $p \leq 0.05$ and $\geq$ 1-fold change; n = 96) represent DEGs enriched in BV2-TurboID pulldown transcriptomes. Black symbols (two-sided t-test; $p \leq 0.05$ and $\geq$

1-fold change; n = 177) represent DEGs enriched in BV2-TurboID bulk transcriptomes. Highly DEGs at the bulk level are mitochondrial-encoded genes. **b.** Volcano plot representation of DEGs between LPS-treated BV2-TurboID cells at the pulldown and bulk level (n=3/group). Green symbols (two-sided t-test; $p \leq 0.05$ and $\geq$ 1-fold change; n = 60) represent DEGs enriched in BV2-TurboID pulldown transcriptomes. Black symbols (two-sided t-test; $p \leq 0.05$ and $\geq$ 1-fold change; n = 108) represent DEGs enriched in LPS-treated BV2-TurboID bulk transcriptomes. Highly DEGs at the LPS-treated bulk level are mitochondrial-encoded genes. **c.** Transcriptome GO analysis of DEGs visualized in panel a. of overrepresented ontologies of non-LPS treated BV2-TurboID cells at the pulldown (left) and bulk (right) levels. **d.** Transcriptome GO analysis of DEGs visualized in panel b. of overrepresented ontologies of LPS-treated BV2-TurboID cells at the pulldown (left) and bulk (right) levels.

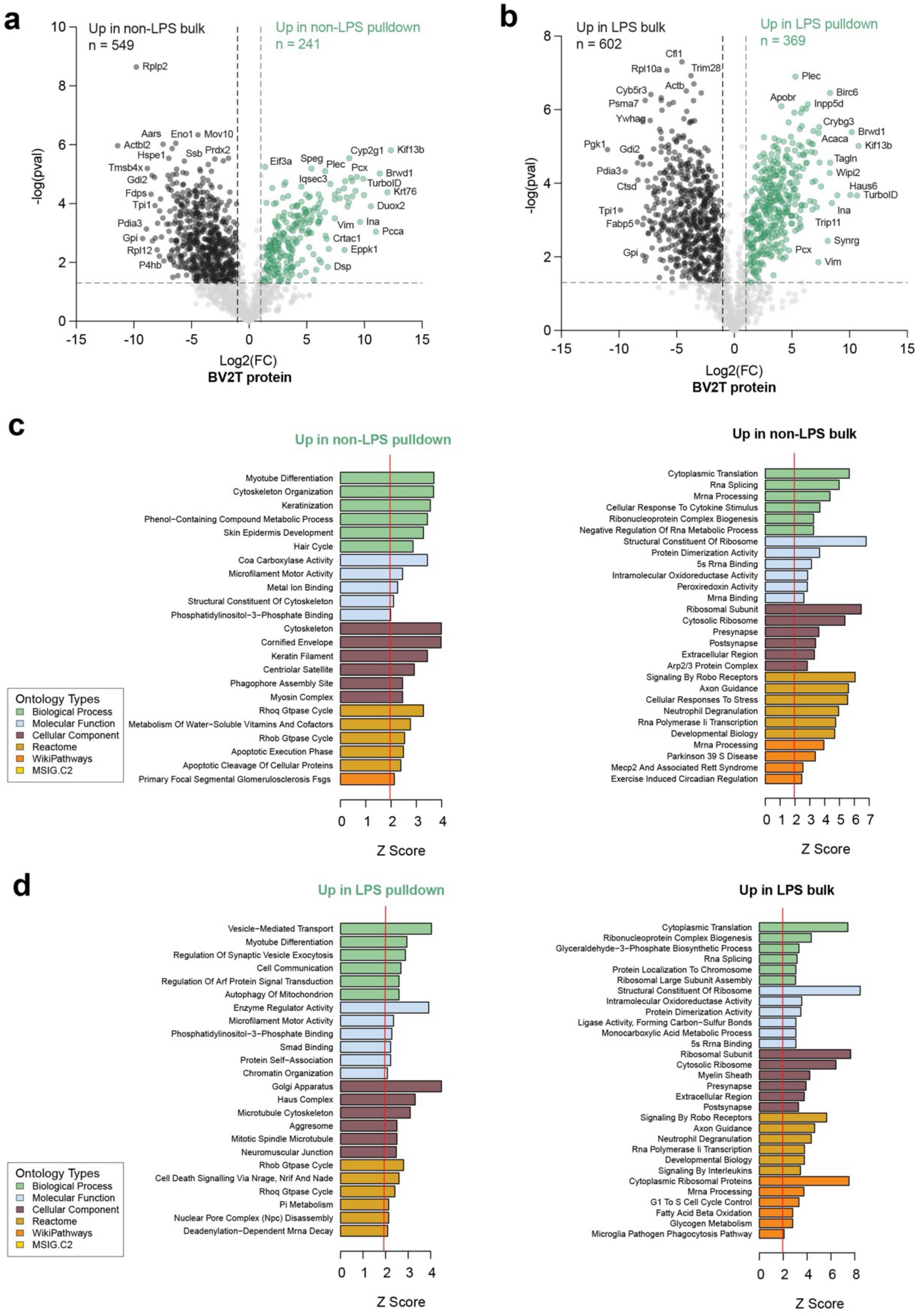


**Supplementary Figure 3: In vitro validation of the SPARO-enriched proteome in untreated and LPS-stimulated BV2-microglial cells.**

**a.** Volcano plot representation of DEPs between non-LPS treated BV2-TurboID cells at the pulldown and bulk level (n=3/group). Green symbols (two-sided t-test; $p \leq 0.05$ and ≥ 1-fold change; n = 241) represent DEPs enriched in BV2-TurboID pulldown proteomes. Black symbols (two-sided t-test; $p \leq 0.05$ and ≥ 1-fold change; n = 549) represent DEPs enriched in BV2-TurboID bulk proteomes. **b.** Volcano plot representation of DEPs between LPS-treated BV2-TurboID cells at the pulldown and bulk level (n=3/group). Green symbols (two-sided t-test; $p \leq 0.05$ and ≥ 1-fold change; n = 369) represent DEPs enriched in BV2-TurboID pulldown proteomes. Black symbols (two-sided t-test; $p \leq 0.05$ and ≥ 1-fold change; n = 602) represent DEGs enriched in LPS-treated BV2-TurboID bulk proteomes. **c.** Proteome GO analysis of DEPs visualized in panel a. of overrepresented ontologies of non-LPS treated BV2-TurboID cells at the pulldown (left) and bulk (right) levels. **d.** Proteome GO analysis of DEPs visualized in panel b. of overrepresented ontologies of LPS-treated BV2-TurboID cells at the pulldown (left) and bulk (right) levels.

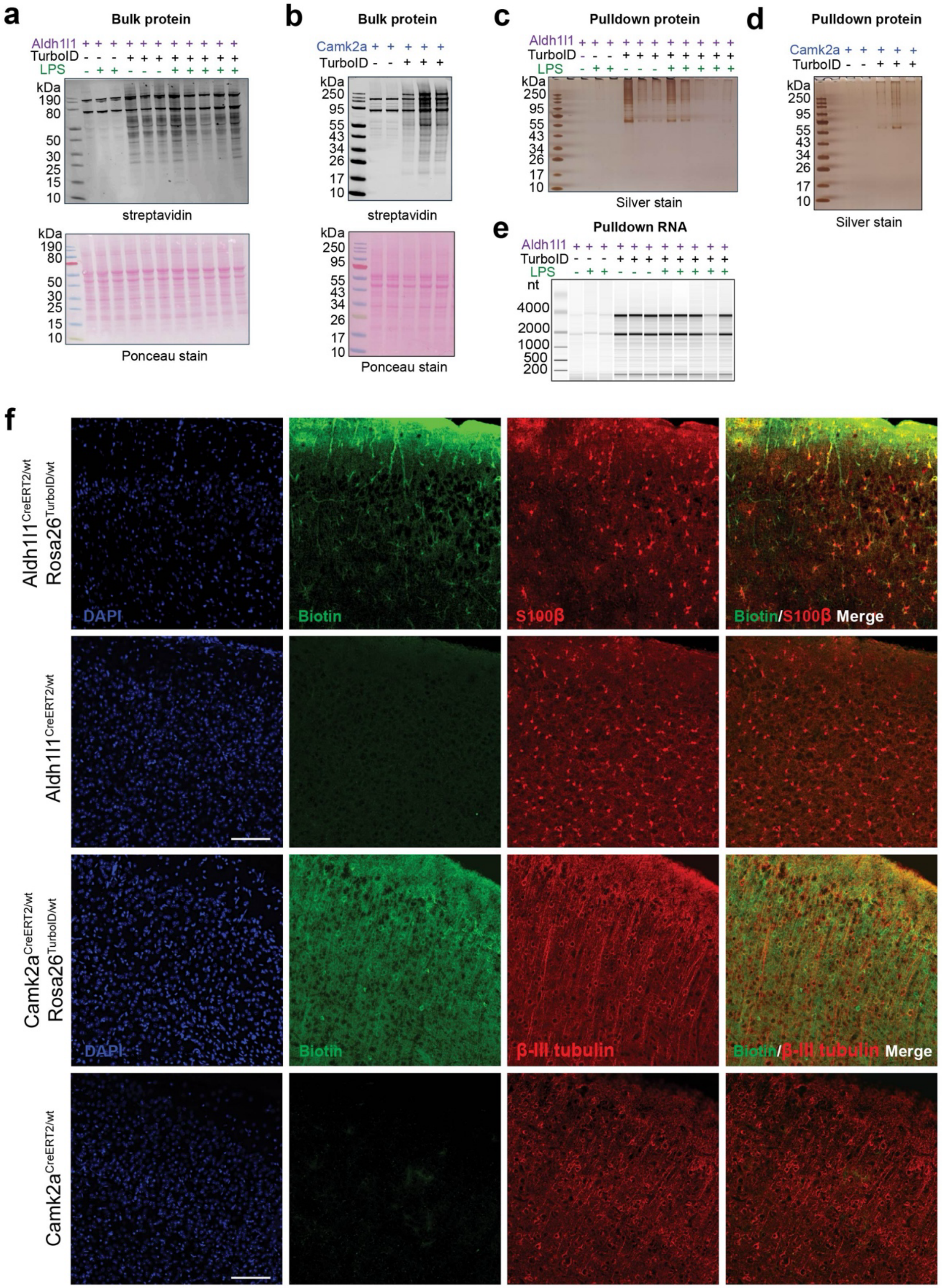


**Supplementary Figure 4: In vivo validation of SPARO in astrocyte-TurboID and neuron-TurboID models.**

**a.** Immunoblot visualization of biotinylated proteins from astrocyte-TurboID bulk cortical lysates, confirming the presence of many biotinylated proteins in the TurboID-expressing animals and only endogenously biotinylated proteins are present in the non-TurboID controls (top, probed with streptavidin-680). Ponceau staining (bottom) of protein abundance showing equal protein loading for immunoblot analysis. **b.** Immunoblot visualization of biotinylated proteins from neuron-TurboID bulk cortical lysates, confirming the presence of many biotinylated proteins in the TurboID-expressing animals and only endogenously biotinylated proteins are present in the non-TurboID controls (top, probed with streptavidin-680). Ponceau staining (bottom) of protein abundance showing equal protein loading for immunoblot analysis. **c.** Silver stain visualization of enriched biotinylated proteins from astrocyte-TurboID cortical lysates after affinity purification from a 10% aliquot of the streptavidin beads. Minimal protein enrichment was found in the control samples, and several proteins were enriched in the astrocyte-TurboID. **d.** Silver stain visualization of enriched biotinylated proteins from neuron-TurboID cortical lysates after affinity purification from a 10% aliquot of the streptavidin beads. Minimal protein enrichment was found in the control samples, and several proteins were enriched in the neuron-TurboID. **e.** Complete RNA gel electrophoresis of RNA eluted off biotinylated proteins after streptavidin pulldown from Fig. 2 panel d. High levels of rRNA and mRNA were detected in astrocyte-TurboID cortical tissue, not seen in non-TurboID controls. **f.** Representative immunofluorescence images of Aldh1l1$^{CreERT2/wt}$ or Camk2a$^{CreERT2/wt}$ controls, astrocyte-TurboID and neuron-TurboID mice sagittal brain sections (n = 3 mice/experimental group) confirms astrocyte and neuron specific biotinylation via streptavidin-488 (green) overlapping with astrocytic marker S100b (red) and neuronal marker b-III tubulin (red), respectively. Nuclei were labeled with DAPI (blue). Scalebar = 100 μm

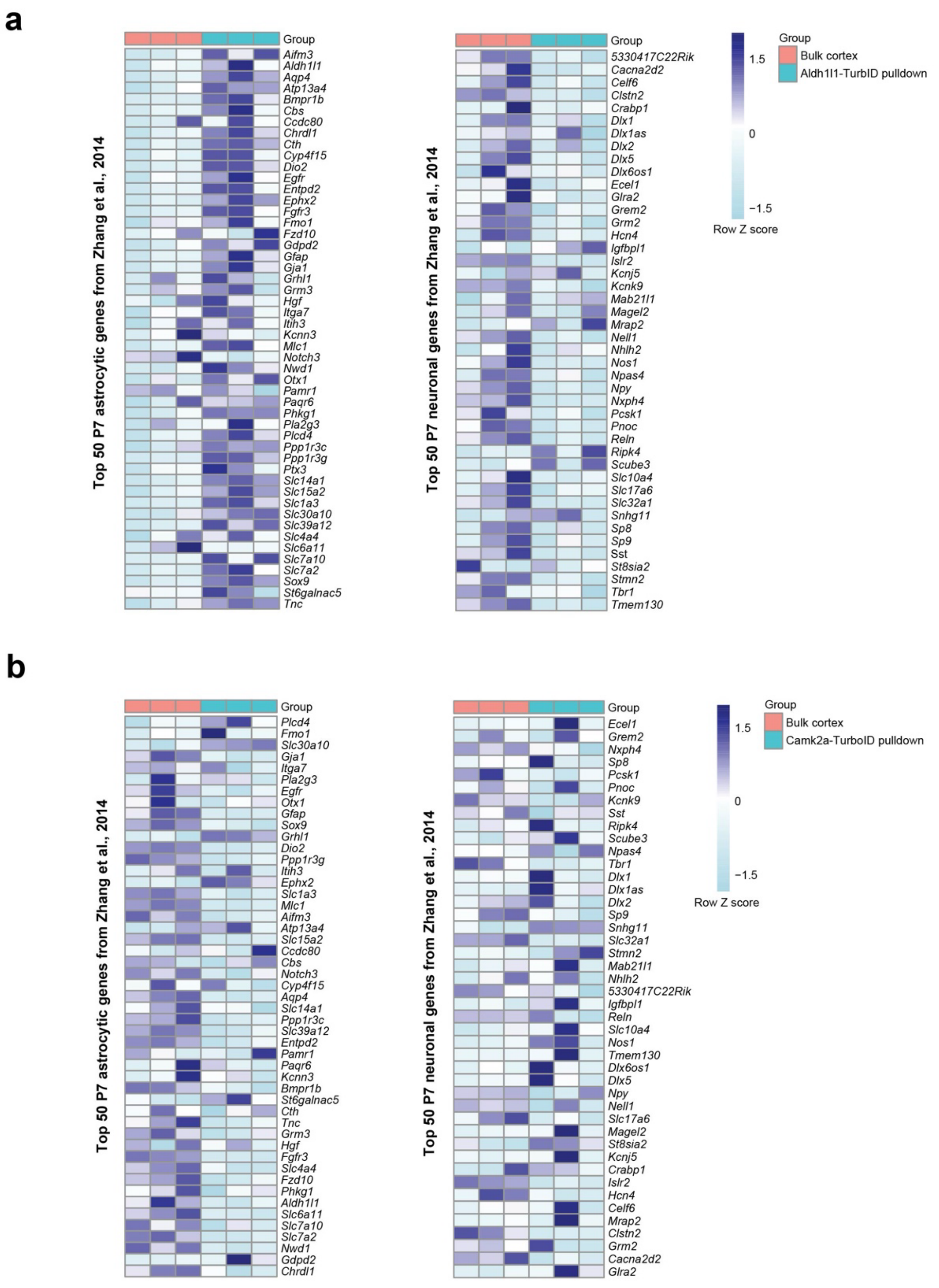


**Supplementary Figure 5: In vivo validation of SPARO to capture cell type enriched pulldown transcriptomes from astrocyte-TurboID and neuron-TurboID models.**

**a.** Heatmap visualization displaying the relative expression (via Row Z score; purple = relative high expression or enrichment and blue = relative low expression or depletion) of the top 50 astrocytic genes (left) and the top 50 neuronal genes (right) present in the bulk and astrocyte-TurboID pulldown samples. The astrocyte-TurboID pulldown samples show a high expression of astrocytic genes and a low expression of neuronal genes. The top 50 astrocyte-specific and neuronal-specific gene lists were acquired from Zhang et al., 2014. **b.** Heatmap visualization displaying the relative expression (via Row Z score; purple = relative high expression or enrichment and blue = relative low expression or depletion) of the top 50 astrocytic genes (left) and the top 50 neuronal genes (right) present in the bulk and neuron-TurboID pulldown samples. The neuron-TurboID pulldown samples show a low expression of astrocytic genes and comparable expression of neuronal genes. The top 50 astrocyte-specific and neuronal-specific gene lists were acquired from Zhang et al., 2014.

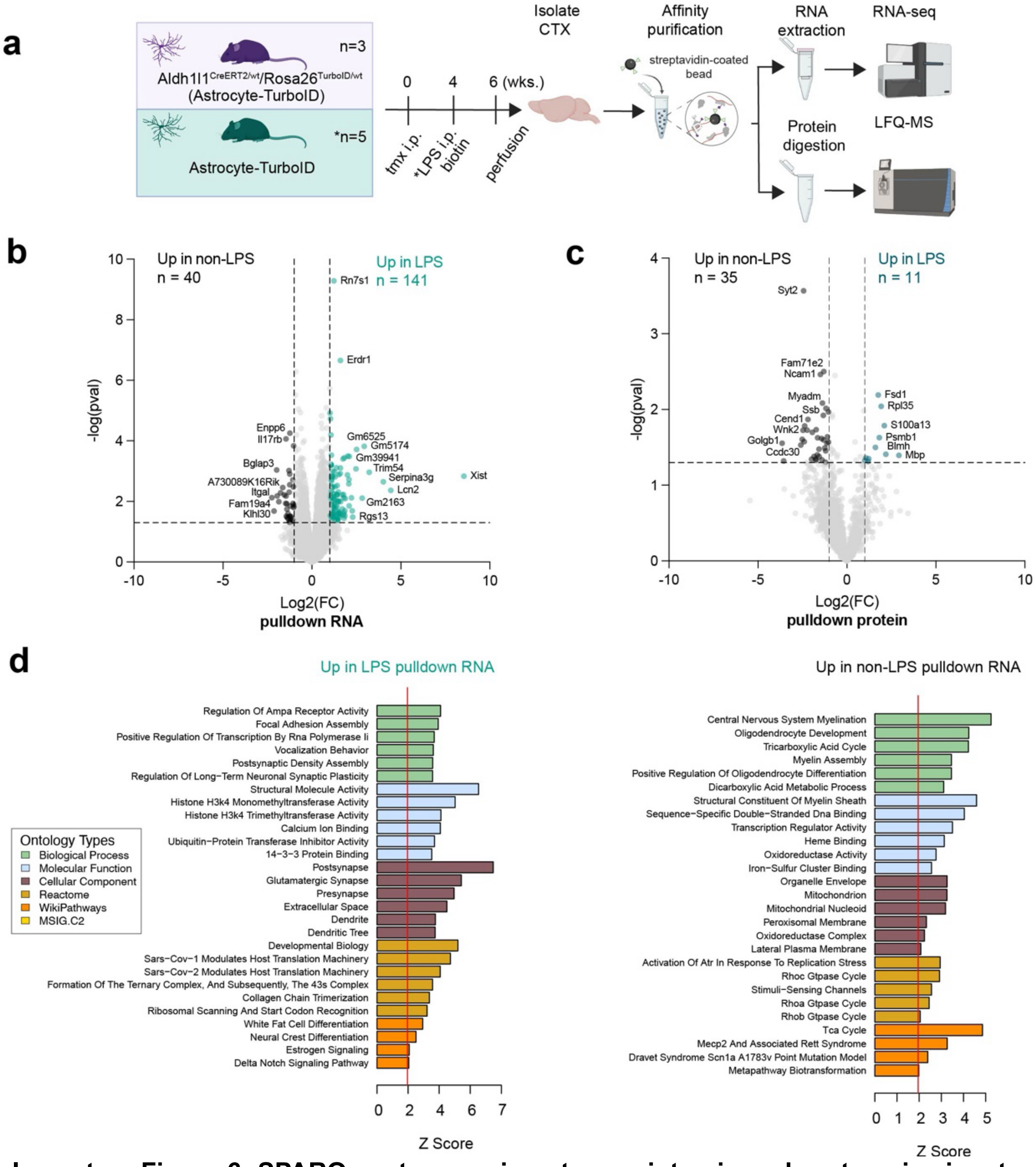

**Supplementary Figure 6: SPARO captures unique transcriptomic and proteomic signature of LPS-stimulated astrocytes in vivo**

**a.** Experimental overview of SPARO in Aldh1l1$^{CreERT2/wt}$/Rosa26$^{TurboID/wt}$ (astrocyte-TurboID, n = 3 mice) and LPS-treated astrocyte-TurboID (n = 5 mice) mouse models. Astrocyte-TurboID mice received tamoxifen (75 mg/kg/day via intraperitoneal (i.p.) injections x 5 days). After 3 weeks of Cre recombination, mice received biotin-containing water (37.5 mg/L) for two weeks. Systemic LPS (0.75 mg/kg/dose x 5 days via i.p. injections) was administered during the first week of biotin water supplementation to induce a neuroinflammatory state. Cortical tissue was isolated for streptavidin affinity purification, followed by LFQ-MS and RNA-sequencing. **b.** Volcano plot representation of DEGs between untreated and LPS-treated astrocyte-TurboID pulldowns. Teal symbols (two-sided t-test; p ≤ 0.05 and ≥ 1-fold change; n = 141) represent pulldown genes enriched in LPS-treated astrocyte-TurboID cortex. Black symbols (two-sided t-test; p ≤ 0.05 and ≥ 1-fold change; n = 40) represent pulldown genes enriched in untreated astrocyte-TurboID cortex. **c.** Volcano plot representation of DEPs between untreated and LPS-treated astrocyte-TurboID pulldowns. Teal symbols (two-sided t-test; p ≤ 0.05 and ≥ 1-fold change; n = 11) represent pulldown proteins enriched in LPS-treated astrocyte-TurboID cortex. Black symbols (two-sided t-test; p ≤ 0.05 and ≥

1-fold change; n = 35) represent pulldown proteins enriched in untreated astrocyte-TurboID cortex. **d.** Transcriptome GO analysis of DEGs visualized in panel b. of overrepresented ontologies of untreated and LPS-treated astrocyte-TurboID pulldowns.

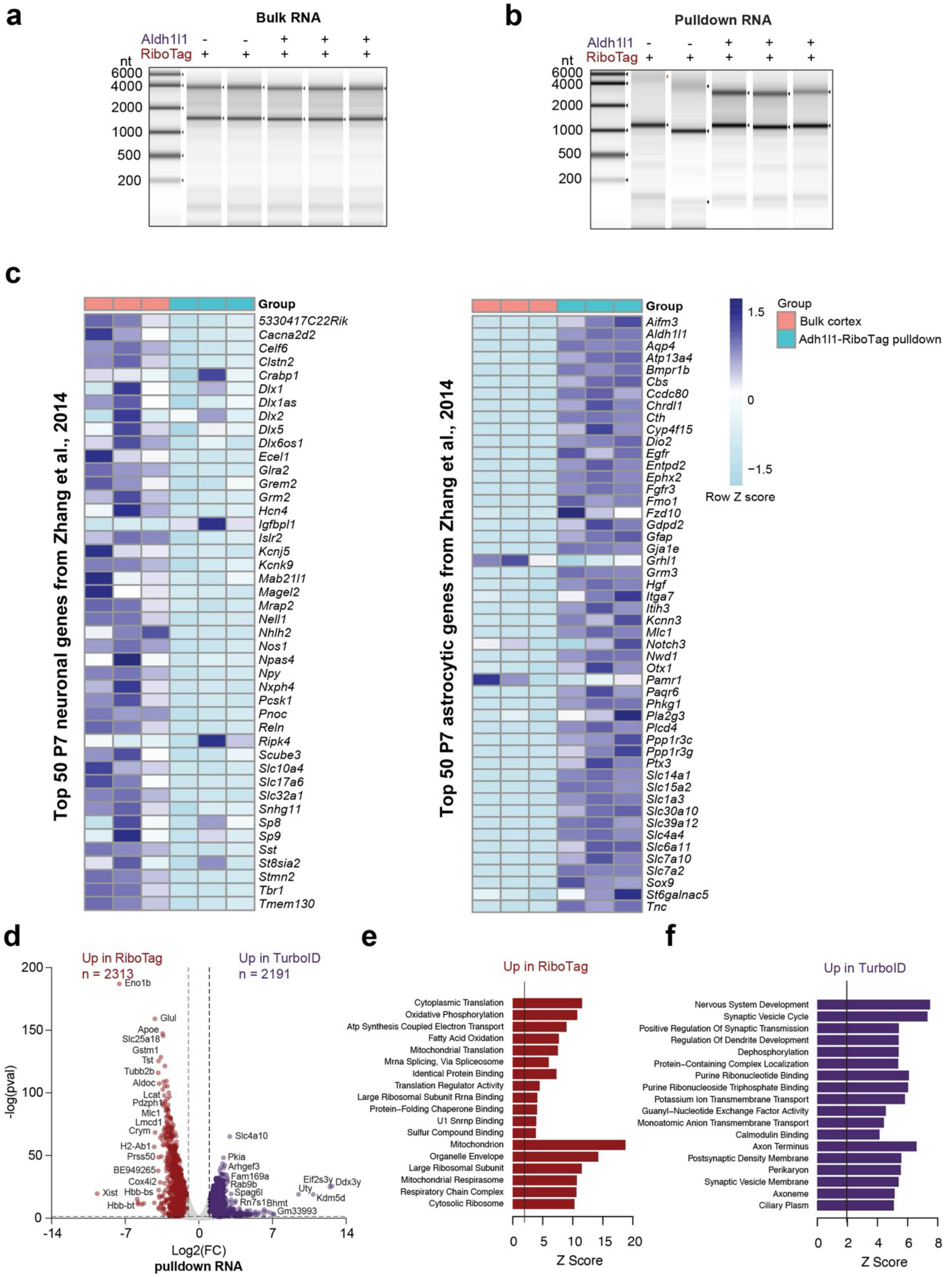


**Supplementary Figure 7: In vivo validation of the astrocyte-RiboTag pulldown transcriptomes and differences with the SPARO enriched astrocyte-TurboID pulldown transcriptomes.**

**a.** RNA gel electrophoresis of high levels of bulk RNA isolated from control and astrocyte-RiboTag animals. **b.** RNA gel electrophoresis of RNA eluted off Rpl22-HA-containing ribosomes after immunoprecipitation. High levels of rRNA and mRNA were detected in astrocyte-RiboTag samples when compared to controls.

**c.** Heatmap visualization displaying the relative expression (via Row Z score; purple = relative high expression or enrichment and blue = relative low expression or depletion) of the top 50 postnatal day 7 (P7) astrocytic genes (left) and the top 50 neuronal genes (right) present in the bulk and astrocyte-RiboTag pulldown samples. The astrocyte-RiboTag pulldown samples show a high expression of astrocytic genes and a low expression of neuronal genes. The top 50 astrocyte-specific and neuronal-specific gene lists were acquired from Zhang et al., 2014. **d.** Volcano plot representation of DEGs between TurboID and RiboTag pulldowns (n=3/group). Red symbols (two-sided t-test; $p \leq 0.05$ and $\geq$ 1-fold change; n = 2313) represent DEGs enriched in RiboTag pulldowns. Purple symbols (two-sided t-test; $p \leq 0.05$ and $\geq$ 1-fold change; n = 2191) represent DEGs enriched in TurboID pulldowns. **e.** Transcriptome GO analysis of DEGs visualized in panel d. of overrepresented ontologies of RiboTag pulldowns including mitochondrial, mRNA processing and translation-related terms. **f.** Transcriptome GO analysis of DEGs visualized in panel d. of overrepresented ontologies of TurboID pulldowns including synaptic, dendritic and ion transport-related terms.

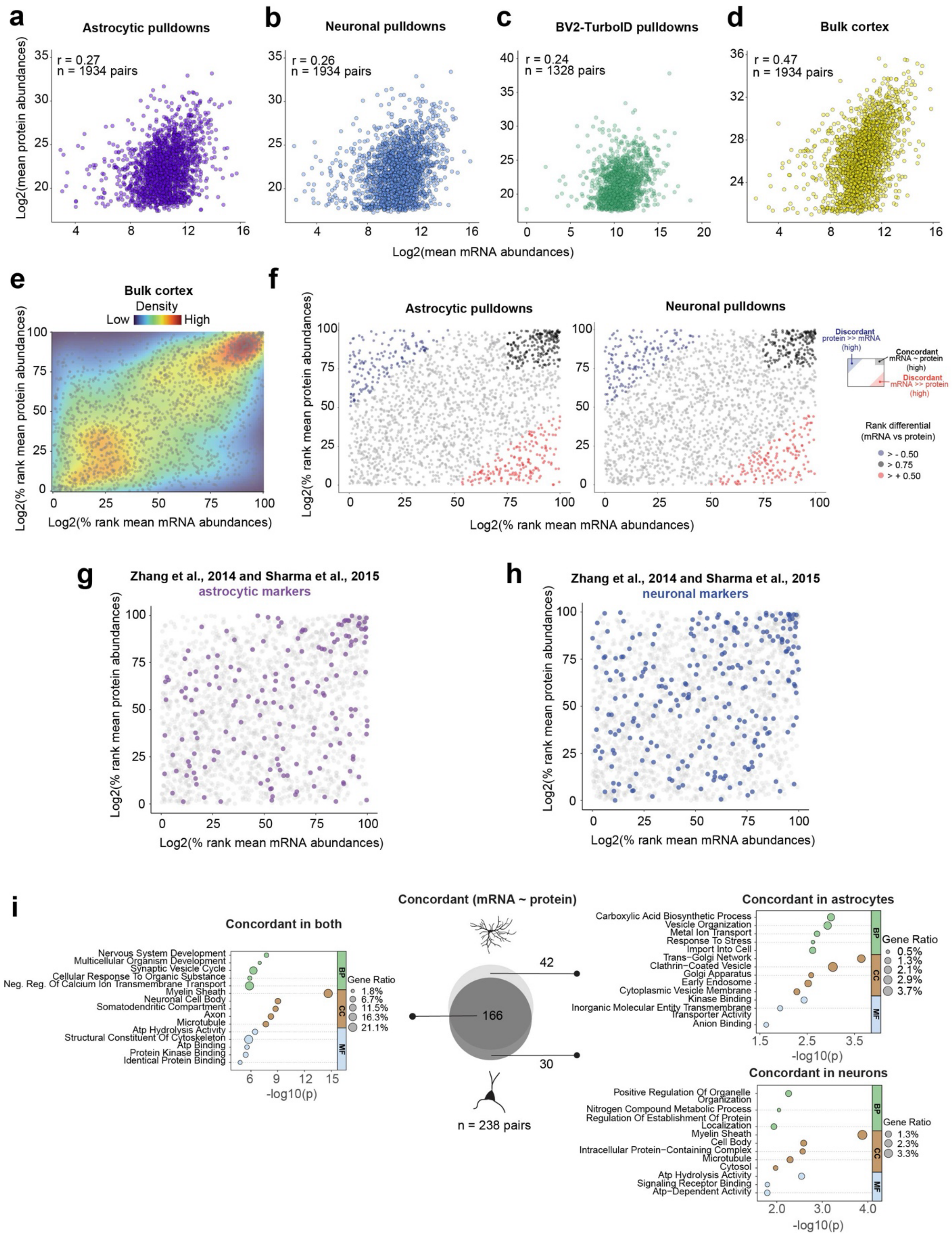
a
Astrocytic pulldowns
r = 0.27
n = 1934 pairs
b
Neuronal pulldowns
r = 0.26
n = 1934 pairs
c
BV2-TurboID pulldowns
r = 0.24
n = 1328 pairs
d
Bulk cortex
r = 0.47
n = 1934 pairs
Log2(mean protein abundances)
Log2(mean mRNA abundances)
e
Bulk cortex
Density
Low
High
Log2(% rank mean protein abundances)
Log2(% rank mean mRNA abundances)
f
Astrocytic pulldowns
Neuronal pulldowns
Discordant protein >> mRNA (high)
Concordant mRNA ~ protein (high)
Discordant mRNA >> protein (high)
Rank differential (mRNA vs protein)
> - 0.50
> 0.75
> + 0.50
g
Zhang et al., 2014 and Sharma et al., 2015
astrocytic markers
h
Zhang et al., 2014 and Sharma et al., 2015
neuronal markers
i
Concordant (mRNA ~ protein)
Concordant in both
Nervous System Development
Multicellular Organism Development
Synaptic Vesicle Cycle
Cellular Response To Organic Substance
Neg. Reg. Of Calcium Ion Transmembrane Transport
Myelin Sheath
Neuronal Cell Body
Somatodendritic Compartment
Axon
Microtubule
Atp Hydrolysis Activity
Structural Constituent Of Cytoskeleton
Atp Binding
Protein Kinase Binding
Identical Protein Binding
Gene Ratio
1.8%
6.7%
11.5%
16.3%
21.1%
-log10(p)
BP
CC
MF
42
166
30
n = 238 pairs
Concordant in astrocytes
Carboxylic Acid Biosynthetic Process
Vesicle Organization
Metal Ion Transport
Response To Stress
Import Into Cell
Trans−Golgi Network
Clathrin−Coated Vesicle
Golgi Apparatus
Early Endosome
Cytoplasmic Vesicle Membrane
Kinase Binding
Inorganic Molecular Entity Transmembrane Transporter Activity
Anion Binding
Gene Ratio
0.5%
1.3%
2.1%
2.9%
3.7%
Concordant in neurons
Positive Regulation Of Organelle Organization
Nitrogen Compound Metabolic Process
Regulation Of Establishment Of Protein Localization
Myelin Sheath
Cell Body
Intracellular Protein−Containing Complex
Microtubule
Cytosol
Atp Hydrolysis Activity
Signaling Receptor Binding
Atp−Dependent Activity
Gene Ratio
1.3%
2.3%
3.3%

**Supplementary Figure 8: Correlation analysis of SPARO transcriptomes and proteomes in vitro and in vivo**

**a.** Correlation analysis between the astrocyte-TurboID pulldown transcriptome (x-axis; via normalized RNA-seq $log_2$-transformed mean count values) and pulldown proteome (y-axis; via normalized LFQ-MS $log_2$-transformed mean intensity values) (n = 1934 gene and protein pairs; Pearson correlation coefficient r = 0.27). **b.** Correlation analysis between the neuron-TurboID pulldown transcriptome (x-axis; via normalized RNA-seq $log_2$-transformed mean count values) and pulldown proteome (y-axis; via normalized LFQ-MS $log_2$-transformed mean intensity values) (n = 1934 gene and protein pairs; Pearson correlation coefficient r = 0.26). **c.** Correlation analysis between the BV2-TurboID pulldown transcriptome (x-axis; via normalized RNA-seq $log_2$-transformed mean count values) and pulldown proteome (y-axis; via normalized LFQ-MS $log_2$-transformed mean intensity values) (n = 1328 gene and protein pairs; Pearson correlation coefficient r = 0.24). **d.** Correlation analysis between the bulk mouse cortex transcriptome (x-axis; via normalized RNA-seq $log_2$-transformed mean count values) and pulldown proteome (y-axis; via normalized LFQ-MS $log_2$-transformed mean intensity values) (n = 1934 gene and protein pairs; Pearson correlation coefficient r = 0.47). **e.** Percent rank transformation and density plot visualization of 1934 gene and protein pairs between the of the bulk mouse cortex (x-axis; via normalized, $log_2$-transformed and percent ranked RNA-seq mean count values) and proteome (y-axis; via normalized, $log_2$-transformed and percent ranked LFQ-MS mean intensity values). **f.** Scatter plot visualization of astrocyte-TurboID (left) and neuron-TurboID (right) pulldown transcriptomes and proteomes (x-axis; via normalized, $log_2$-transformed and percent ranked RNA-seq mean count values) and proteome (y-axis; via normalized, $log_2$-transformed and percent ranked LFQ-MS mean intensity values). The highly concordant (mRNA ~ protein in black) or discordant (protein >> mRNA in blue or mRNA >> protein in red) groups were nominated using a rank differential value (mRNA – protein percentile abundances). **g.** Scatter plot visualization of the data points in panel f (left) with astrocytic markers color-coded in purple. The astrocyte-specific marker list was acquired from the unionization of top astrocytic markers from Zhang et al., 2014 and Sharma et al., 2015. **h.** Scatter plot visualization of the data points in panel f (right) with neuronal markers color-coded in blue. The neuron-specific marker list was acquired from the unionization of top neuronal markers from Zhang et al., 2014 and Sharma et al., 2015. **i.** Venn diagram presentation of mRNA and protein pair quantities present in the concordant (top right; black; mRNA ~ protein) group between astrocyte-TurboID and neuron-TurboID paired pulldown transcriptomes and proteomes.

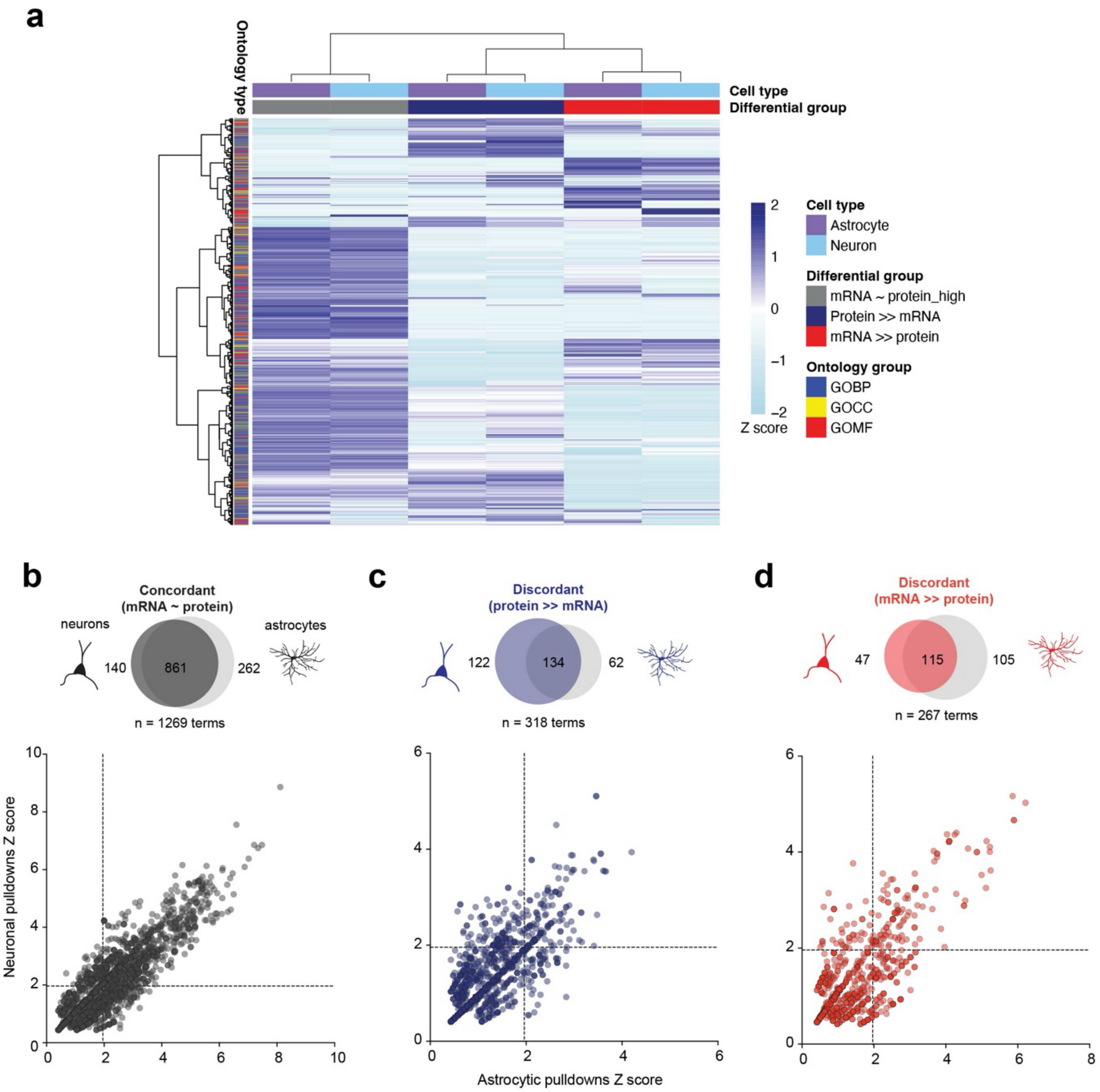


**Supplementary Figure 9: GO term overlap of concordant and discordant mRNA-protein pairs of astrocyte-TurboID and neuron-TurboID SPARO pulldowns.**

**a.** Heatmap visualization of GO terms (via Z score enrichment) between astrocyte-TurboID and neuron-TurboID paired pulldown transcriptomes and proteomes from each concordant (mRNA ~ protein in black) and discordant (protein >> mRNA in blue or mRNA >> protein in red) groups. **b.** Venn diagram presentation (top) of GO term quantities present in the concordant (top right; black; mRNA ~ protein) group between astrocyte-TurboID and neuron-TurboID paired pulldown transcriptomes and proteomes. In the neuronal pulldowns, n = 140 terms and in the astrocytic pulldowns n = 262 terms were unique with n = 861 terms overlapping between the two cell types. Scatter plot visualization (bottom) of GO term Z scores from astrocytic pulldowns (x-axis) and neuronal pulldowns (y-axis). **c.** Venn diagram presentation (top) of GO term quantities present in the discordant (top left; blue; protein >> mRNA) group between astrocyte-TurboID and neuron-TurboID paired pulldown transcriptomes and proteomes. Scatter plot visualization (bottom) of GO term Z scores from astrocytic pulldowns (x-axis) and neuronal pulldowns (y-axis). **d.** Venn diagram presentation (top) of GO term quantities present in the discordant (bottom right; red; mRNA >> protein) group between astrocyte-TurboID and neuron-TurboID paired pulldown transcriptomes and proteomes. Scatter plot visualization (bottom) of GO term Z scores from astrocytic pulldowns (x-axis) and neuronal pulldowns (y-axis).

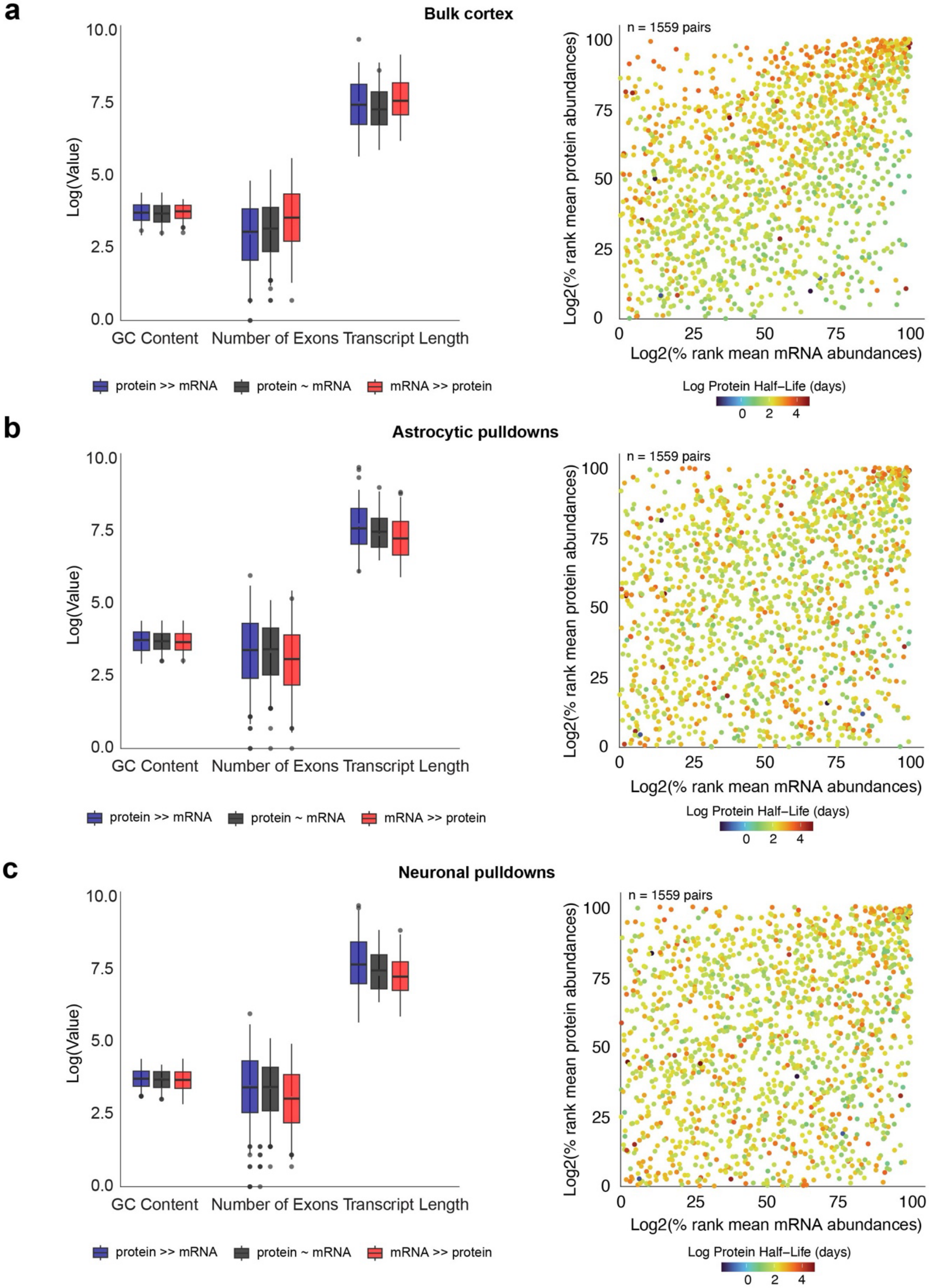

a
Bulk cortex
Log(Value)
GC Content
Number of Exons
Transcript Length
protein >> mRNA
protein ~ mRNA
mRNA >> protein
n = 1559 pairs
Log2(% rank mean protein abundances)
Log2(% rank mean mRNA abundances)
Log Protein Half-Life (days)
b
Astrocytic pulldowns
Log(Value)
GC Content
Number of Exons
Transcript Length
protein >> mRNA
protein ~ mRNA
mRNA >> protein
n = 1559 pairs
Log2(% rank mean protein abundances)
Log2(% rank mean mRNA abundances)
Log Protein Half-Life (days)
c
Neuronal pulldowns
Log(Value)
GC Content
Number of Exons
Transcript Length
protein >> mRNA
protein ~ mRNA
mRNA >> protein
n = 1559 pairs
Log2(% rank mean protein abundances)
Log2(% rank mean mRNA abundances)
Log Protein Half-Life (days)

**Supplementary Figure 10: Investigating transcript traits and protein half-life to identify processes contributing to discordance between mRNA and protein abundances.**

**a.** Boxplot visualization of RNA-seq transcript traits: average GC content (%), number of exons and the average transcript length (nucleotides) for the bulk cortex concordant (mRNA ~ protein in black) and discordant (protein >> mRNA in blue or mRNA >> protein in red) groups (left) and the distribution of the log of protein half-life in days (right). RNA trait values are derived from the BioMart/ENSEMBL databases. Protein half-life values are derived from Fornasiero et al., 2018. **b.** Boxplot visualization of RNA-seq transcript RNA traits: average GC content (%), number of exons and the average transcript length (nucleotides) for the astrocyte-TurboID pulldown concordant (mRNA ~ protein in black) and discordant (protein >> mRNA in blue or mRNA >> protein in red) groups (left) and the distribution of the log of protein half-life in days (right). **c.** Boxplot visualization of RNA-seq transcript RNA traits: average GC content (%), number of exons and the average transcript length (nucleotides) for the neuron-TurboID pulldown concordant (mRNA ~ protein in black) and discordant (protein >> mRNA in blue or mRNA >> protein in red) groups (left) and the distribution of the log of protein half-life in days (right).
